# Concerted actions of distinct serotonin neurons orchestrate female pup care behavior

**DOI:** 10.1101/2025.07.31.667987

**Authors:** Shuyun Alina Xiao, Che Cherry Chen, Patricia Horvath, Valerie Tsai, Vibiana Marie Cardenas, Dan Biderman, Fei Deng, Yulong Li, Scott W. Linderman, Catherine Dulac, Liqun Luo

## Abstract

In many mammalian species, female behavior towards infant conspecifics changes across reproductive stages. Sexually naïve females interact minimally or aggressively with infants, whereas the same animals exhibit extensive care behavior, even towards unrelated infants, after parturition^1–6^. Here, we discovered that two distinct sets of serotonin neurons collectively mediate this dramatic transition in maternal behavior—serotonin neurons projecting to the medial preoptic area (mPOA) promote pup care in mothers, whereas those projecting to the bed nucleus of the stria terminalis (BNST) suppress pup interaction in virgin female mice. Disrupting serotonin synthesis in either of these subpopulations or stimulating either subpopulation is sufficient to toggle pup-directed behavior between that displayed by virgin females and that of lactating mothers. In virgin female mice, the first pup interaction triggers an increase in serotonin release in BNST but a decrease in mPOA. In mothers, serotonin activity becomes greatly elevated in mPOA during pup interactions. Acute interruption of serotonin signaling locally in either mPOA or BNST disrupts the stage-dependent switch in pup care. Together, these results highlight how functionally distinct serotonin subpopulations orchestrate social behaviors appropriate for a given reproductive state, and suggest a circuit logic for how a neuromodulator coordinates adaptive behavioral changes across life stages.

## Main

Social behaviors are dynamic throughout an animal’s lifetime to meet the diverse needs of different life stages. Parental behavior, for instance, is indispensable to the survival of the young in many species^2,3,5^. In mice, naïve virgin females ignore, avoid, and even kill pups^1–5^. However, after parturition, the same animals fundamentally shift their behavioral and physiological priorities^1–5^. They not only care for their own pups but also readily foster nonbiological offspring^6^. How the brain modulates such drastic behavioral switches in a life stage appropriate manner remains elusive.

Serotonin (5-hydroxytryptamine, or 5-HT), a phylogenetically ancient neuromodulator, is necessary for a diverse array of social behaviors^7–13^, including the proper expression of maternal care^14–16^. Depleting brain serotonin by knocking out tryptophan hydroxylase 2 (Tph2), an enzyme required for serotonin biosynthesis in the central nervous system, substantially reduces the quantity and quality of maternal care in lactating mice^14^. However, how serotonin functions at the circuit level to modulate care behavior across reproductive stages is unknown. Although small in neuronal number and anatomically restricted to the raphe nuclei, serotonin neurons comprise one of the largest efferent systems^17^ and contain subpopulations characterized by distinct projection patterns, physiology, and functions^18–21^. While this complex anatomy has left many of serotonin’s functions unknown at the fine neural circuit level, the exact extensiveness of its neural architecture hints at serotonin’s rich potential for broadcasting information across the brain and coordinate behavioral switches in the same animal.

Here, we systematically mapped the axons of serotonin neurons activated by pup interactions and tested the role of serotonin across projections, social contexts, and reproductive states in female mice. We discovered two functionally and physiologically distinct serotoninergic subpopulations that together orchestrate the state-dependent change in female pup care behavior.

### Pup interaction activates 5-HT neurons

To assess which social interactions activate serotonin neurons, we tracked each female mouse across reproductive states and queried serotonin neuron activity using population Ca^2+^ recordings during a wide gamut of social behaviors, including interactions with a juvenile female, interactions with pups both as a virgin and as a lactating mother, as well as sexual interactions during different phases of her estrus cycle (Fig. 1a, d). We used double transgenic mice that carry Cre recombinase in cells expressing serotonin transporter (*Sert-Cre*)^22^ and a Cre- dependent Ca^2+^ indicator reporter (*Ai148*)^23^ for stable, consistent GCaMP6f expression across serotonin neurons. We implanted an optic fiber over either dorsal or median raphe (DR or MR) (Fig. 1a), where cell bodies of serotonin neurons that innervate the forebrain and midbrain reside^17^. Over 98% of GCaMP6f^+^ neurons in the raphe co-expressed the serotonin marker Tph2, while 75% of raphe serotonin neurons expressed GCaMP6f, validating the specificity and efficiency of our transgenic approach (Fig. 1b, c).

**Fig. 1.**
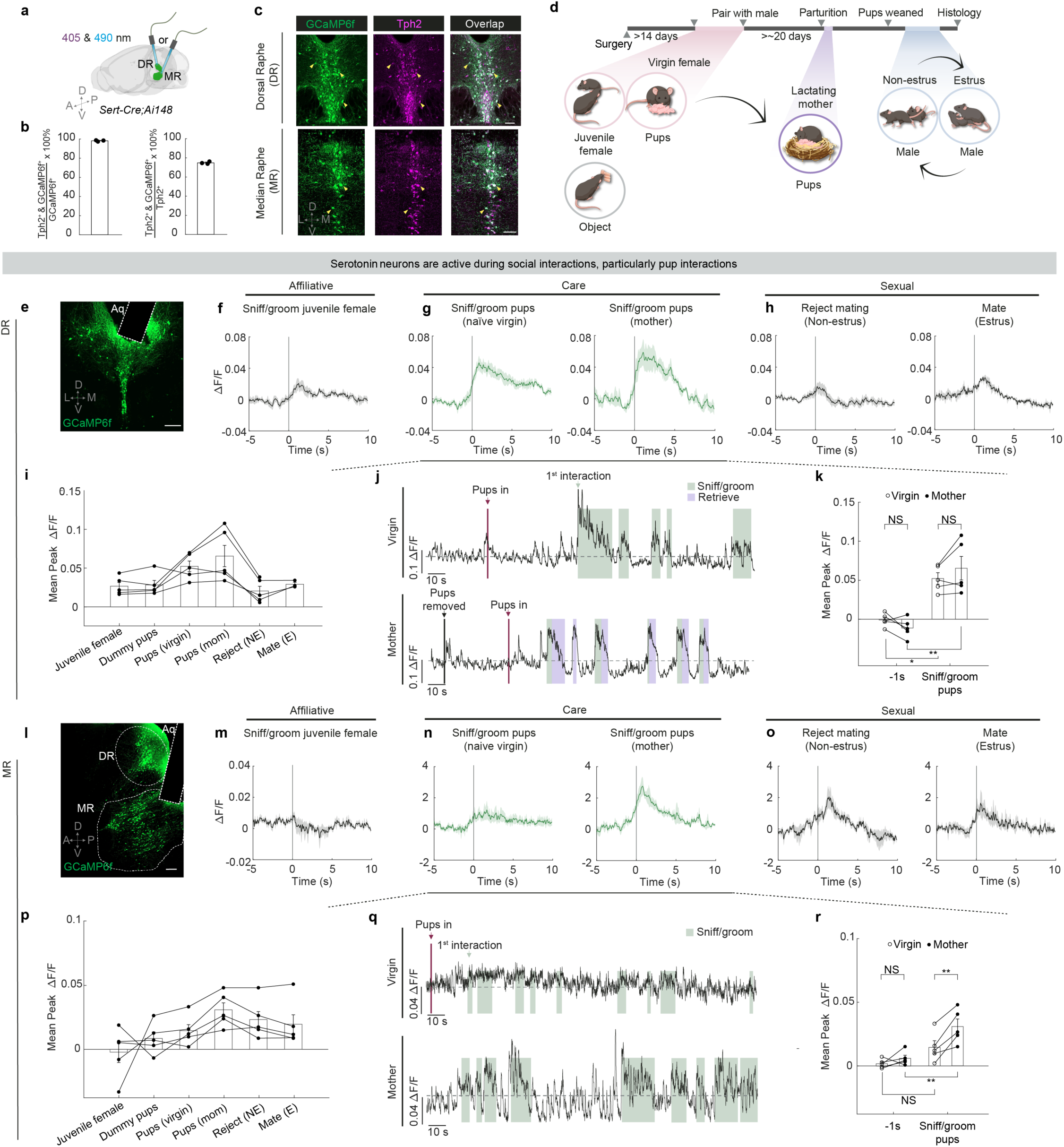
Serotonin neurons are active during social interactions. **a**, Schematic of fiber photometry recording of serotonin neurons in DR and MR. All neural signals were recorded at two wavelengths: 490 nm that captures Ca^2+^-dependent fluorescence and 405 nm, an isosbestic control that captures Ca^2+^-independent background noise and motion artifacts. **b**, Percentage of GCaMP6f cells co-expressing Tph2 (left) and percentage of Tph2 cells co-expressing GCaMP6f in the raphe (right). *n* = 3 mice. **c**, Representative confocal images showing the overlap of GCaMP6f (green) and Tph2 (magenta) in *Sert- Cre;Ai148* mice. Arrowheads point to example double-labeled cells. Scale bars, 100 μm. **d**, Experimental timeline for the six behaviors during which photometry recording was performed in each female. Black arrows highlight transitions in female reproductive states. **e,** Representative histology showing GCaMP6f expression (green) and fiber track (dotted white line) in DR, coronal view. Scale bar, 200 μm. **f–h**, Peri-event time histograms (PETHs) of Δ*F*/*F* (relative change in the 490-nm fluorescence signal after removing baseline fluctuations estimated from the 405-nm control signal) of DR serotonin neurons aligned to the onset of females sniffing/grooming a juvenile female (**f**), sniffing/grooming pups as naive virgins (**g**, left) and as mothers (**g**, right), rejecting male mounting during non-estrus (**h**, left), and mating upon male mounting during estrus (**h**, right) (*n* = 5 for **f**–**h**, left, *n* = 3 for **h**, right). **i**, Mean peak Δ*F*/*F* of DR serotonin neurons during the six behaviors (**d**), with lines linking data from the same mice. **j**, Representative Δ*F*/*F* traces of DR serotonin neurons during pup interactions in the same female as a naive virgin (**j**, top) and a mother (**j**, bottom). Gray dash line indicates Δ*F*/*F* = 0. **k**, Responses before (–1 sec relative to behavior onset) and during pup sniffing/grooming across reproductive stages [*n* = 5; mixed-effects model: Restricted Maximum Likelihood (REML)–fixed effects type III followed by Šídák’s multiple comparisons test]. **l,** Same as (**e**) but in MR, sagittal view. **m–o**, Same as **f–h**, but in MR (*n* = 5). **p–r**, Same as **i–k**, but in MR. Data are shown as mean ± s.e.m. Abbreviation in this and all subsequent figures: DR, dorsal raphe; MR, median raphe. A, anterior; P, posterior; D, dorsal; V, ventral; L, lateral; M, medial. * *p* < 0.05; ** *p* < 0.01; *** *p* < 0.001; NS, not significant. See Source Data for detailed statistics.

Serotonin neurons, especially those we recorded in the DR, were active during interactions with same-sex conspecifics and aspects of mating behaviors (Fig. 1f, h, i, m, o, p; Extended Data Fig. 1), consistent with previous studies^9,10,24–26^. However, among the behaviors recorded, we found that serotonin neurons across the two raphe locations were most robustly activated during pup interactions in mothers (Fig. 1 e–i, l–p; Extended Data Fig. 1e, j); yet, they had different response patterns in virgins.

In DR, Ca^2+^ signal increased strongly upon the first pup encounter in virgins, and showed recurring increases, albeit weaker than the first response, during subsequent interactions (Fig. 1g, j; Extended Data Fig. 1k). These pup-directed increases in Ca^2+^ signal were stronger and more sustained than those during dummy “pup” investigation despite similarity in interaction time (Extended Data Fig. 1n), indicating that the heightened activation by pups was unlikely a generic response to novel small objects. After these female mice became mothers, activity of DR serotonin neurons also increased strongly when the recorded animals sniffed and groomed pups (Fig. 1g, j). Nonetheless, response magnitudes during these pup interactions did not differ significantly between virgins and mothers (Fig. 1k).

By contrast, MR serotonin neurons responded weakly to both real and dummy pups in virgins (Fig. 1n, q; Extended Data Fig. 1o, r). However, after parturition, MR serotonin neurons became strongly activated when they sniffed and groomed the pups (Fig. 1n, q, r). Together, the differences in pup-induced response across anatomical locations and reproductive stages reveal that the physiology of serotonin neurons not only is heterogeneous but also can be dynamic across an animal’s lifetime. This observation suggests that the serotonin system might play a more nuanced role than to invariably promote pup interaction.

### Mapping active 5-HT projections

Serotonin neurons comprise heterogenous populations with distinct projection targets^18,19,21,27^. To identify projection patterns of serotonin neurons activated by pup interaction comprehensively across the entire brain, we intersected *Sert-Flp* transgenic mice^19^ with the TRAP2 system^28^, which captures pup-activated serotonin neurons at high efficiency and specificity (Extended Data Fig. 2a–d). In virgin females, we injected an AAV that expresses mCherry doubly gated by Cre and Flp (Cre-On, Flp-On) into the DR and MR (Fig. 2a). Two weeks later, we administered 4-hydroxytamoxifen (4OHT) following one hour of either pup interaction in the home cage (pup- TRAP) or home cage activity alone (HC-TRAP). Taking advantage of the long TRAP window (∼± 3 hours relative to 4OHT administration^28,29^) and the fact that many virgin females begin to exhibit some level of pup care after prolonged pup exposure^30,31^, we let the pup-TRAP females resume pup interactions for another three hours after 4OHT injection to capture a mix of pup avoidance-related and care-related serotoninergic projections in the same animal.

**Fig. 2.**
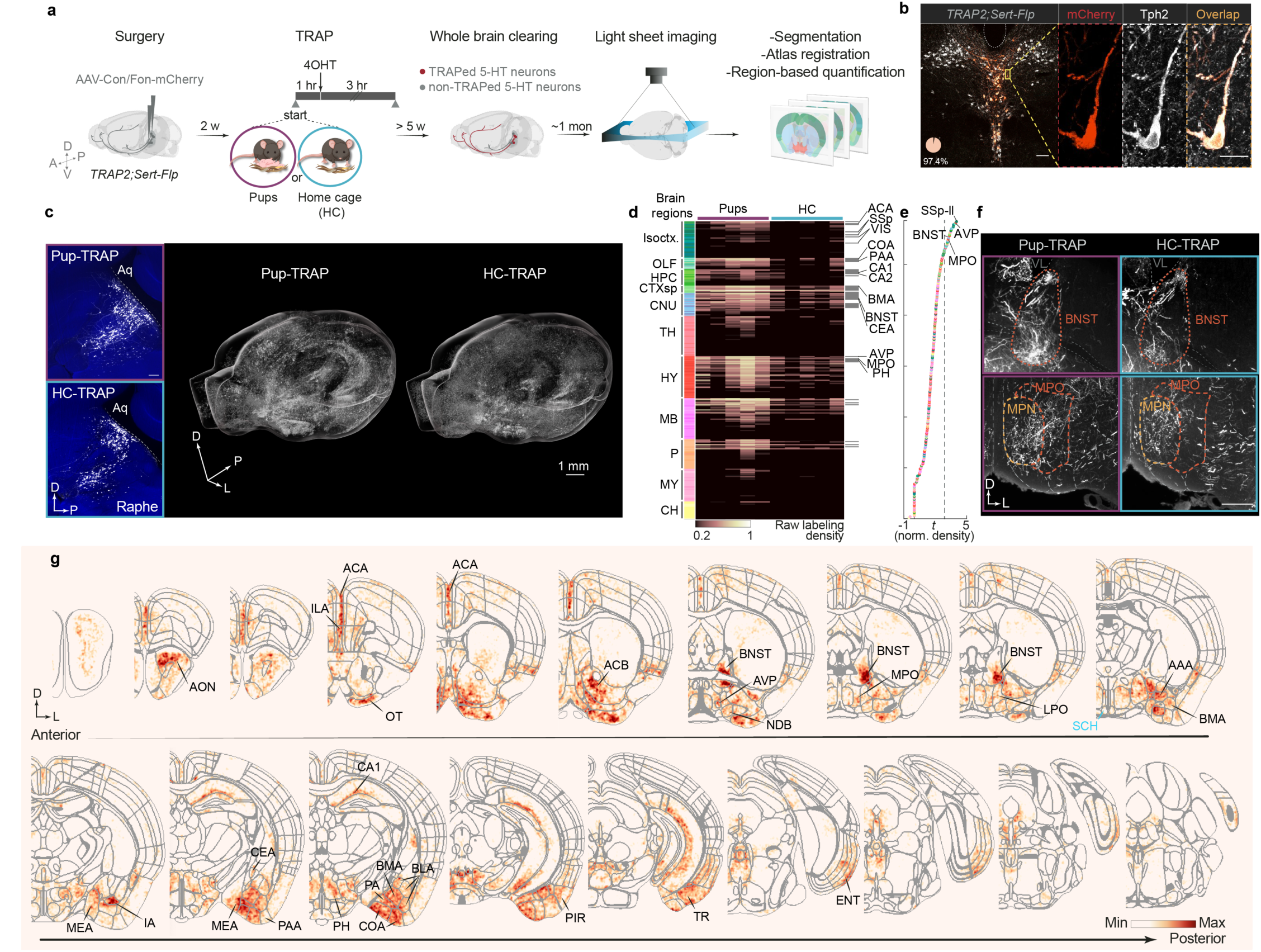
Whole-brain axon mapping of serotonin neurons activated by pup interaction. a, Schematic of viral-genetic strategy and whole-brain imaging. **b**, Representative histology showing the overlap (orange, mCherry^+^ & Tph2^+^ / mCherry^+^ = 97.4% from two mice) of mCherry (red) and Tph2 expression (gray) in the DR of a 4OHT-injected *TRAP2;Sert-Flp* mouse, coronal view. Scale bars, 100 μm (20 μm in enlarged view of neurons outlined in the boxed areas). **c**, Representative histology showing raphe serotonin cell bodies from an example pup-TRAPed brain (magenta box) and home cage (HC)-TRAPed brain (cyan box), sagittal view. Scale bar, 200 μm (**c**, left). 3D rendering of axonal projections from the same two brains (**c**, right). Aq, aqueduct. **d**, Heatmap showing raw labeling density across 292 regions defined by the Allen Atlas (*n* = 5 pup-TRAP, *n* = 5 HC-TRAP, with each column representing a mouse). Gray horizontal lines to the right show findings that pass false discovery rate at 3%. **e**, Difference in range-normalized labeling density between pup-TRAP and HC-TRAP conditions shown by *t*- statistic. Dashed line demarcates the top 30 regions that differ between the two conditions in normalized labeling density. Dots are color coded by brain regions per Allen Brain Atlas coloring convention. **f**, Maximum intensity projection of 60-μm optical sections from the same two representative brains in selected target regions, including BNST (top) and mPOA (which includes both MPO and MPN defined by Allen Brain Atlas, bottom). Sections are shown in coronal view. Scale bars, 500 μm. **g**, Coronal maps showing the voxel-wise difference in average axon labeling density between the pup-TRAPed and HC-TRAPed brains. Darker red indicates higher mean axon density in the pup-TRAP condition. Black lines and text highlight a subset of regions with higher axon labeling density in the pup-TRAP condition. Cyan line and text annotate SCH, a region densely innervated by serotonin neurons but is not differentially labeled between pup- and HC-TRAP conditions. See Supplementary Video 1 for the fly-through of coronal maps showing all brain regions. See Extended Data Fig. 2f for explanations of anatomical region abbreviations, and Source Data for detailed statistics.

Since Cre expression is only induced in neurons activated by the TRAPed experience and Flp is only expressed in *Sert*^+^ neurons, mCherry expression should be restricted to pup-activated or HC-activated serotonin neurons. Indeed, >97% of the neurons expressing mCherry co-expressed Tph2, confirming that the labeled axons stem predominantly from serotonin neurons (Fig. 2b). At least five weeks after TRAP, we dissected the brains and performed whole brain clearing, imaged the brains using light sheet microscopy, segmented the axons using TrailMap^32^, and registered the volumes to the Allen reference brain to compare the axon density between the two TRAP conditions across mice in the same reference space (Fig. 2a). In both HC- and pup-TRAP conditions, we captured serotonin neurons across DR and MR, with widespread axons across the brain (Fig. 2c). Consistent with our observation that pup interactions induced higher Ca^2+^ activity in serotonin neurons, more cells were labeled in the pup- than the HC-TRAP condition overall, with several regions showing markedly higher axon density in the pup-TRAP condition (Fig. 2c–g; Extended Data Fig. 2e; Supplementary Video 1).

Whole-brain region-based quantifications revealed that many of these hotspots were in cortical regions, cerebral nuclei, and hypothalamus (Fig. 2d). Serotoninergic axons innervating amygdala and extended amygdala nuclei such as bed nucleus of the stria terminalis (BNST) were broadly captured and among the highest differentially labeled projections in the pup-TRAP condition (Fig. 2d, f, g; Extended Data Fig. 2f). Various sensory related areas, including olfactory, somatosensory, and visual cortical areas showed consistently more labeling in the pup-TRAPed brains (Fig. 2d). Within the hypothalamus, certain regions such as medial preoptic area (mPOA) showed much denser axon labeling in the pup-TRAP than the HC-TRAP condition (Fig. 2d, f). Notably, many nearby regions such as suprachiasmatic nucleus (SCH) were not significantly different in axon labeling between the two conditions (Fig. 2d, g), even though they are known to receive dense innervation by serotonin neurons^33^. This indicates that pup-TRAP does not indiscriminately label all serotoninergic targets, but instead selectively highlights a suite of regions potentially recruited for social interaction.

To further test if the differences in raw labeling density hinged solely on the fact that more serotonin neurons were labeled in the pup-TRAPed brains, we compared the normalized axon density between the two TRAP conditions. Across the two comparisons—using respectively the raw and normalized axon density—a subset of the same regions including BNST and mPOA remained the top differentially labeled targets in the pup-TRAP condition (Fig. 2e). These two quantification methods support the notion that serotonin neurons activated by pup interaction not only project heavily to these regions but show a spatial preference for them, too.

Previous studies indicate that mPOA and BNST drive opposing functions in pup-directed behaviors through reciprocal inhibitory connections^4,34–36^. Whereas estrogen receptor alpha–expressing (*Esr1^+^*) neurons in the mPOA promote pup care and suppress the activity of *Esr1*^+^ neurons in BNST, the latter antagonize the activity of the former and drive infanticide in certain mouse strains^4^. However, beyond classic hormones such as estradiol and progesterone, which have been shown recently to increase the selectivity of mPOA *Galanin*^+^ neurons to pup stimuli during pregnancy^37^, how these two circuit nodes are modulated in virgins and lactating mothers remains elusive. The recruitment of serotonin axons that project to mPOA and BNST raises the possibility that serotonin might be one source of neuromodulation that shifts the balance between these two key sites to predispose different pup-directed behaviors. We thus focused on mPOA- and BNST-projecting serotonin neurons (hereafter abbreviated as 5HT_mPOA_ and 5HT_BNST_, respectively) in the following experiments.

To test whether the same serotonin neurons innervate both regions, we conducted *Sert-Cre*-gated retrograde tracing to visualize the cell body location of 5HT_mPOA_ and 5HT_BNST_. Although both serotonin neuron subpopulations had cells across DR and MR, 5HT_mPOA_ were preferentially located in the MR and 5HT_BNST_ in the DR (Extended Data Fig. 3a–d). Dual retrograde tracing in the same brains further showed that 5HT_mPOA_ and 5HT_BNST_ exhibited little overlap (Extended Data Fig. 3e–h). Since DR and MR 5-HT neurons have, respectively, different response patterns to pups across life stages (Fig. 1), the labeling of 5HT_mPOA_ and 5HT_BNST_ in the pup- TRAP condition raises the question of whether these serotonin subpopulations themselves are functionally distinct in regulating a female’s response towards pups.

### 5HTmPOA promotes maternal care

We next tested the functional role of 5HT_mPOA_ and 5HT_BNST_ subpopulations. As many serotonin neurons co-release other neurotransmitters and neuropeptides^19,38–40^, we employed a conditional knockout strategy to investigate specifically the function of serotonin. In *Tph2^flox/flox^* mice^41^, we bilaterally injected *AAVretro-Cre-P2A-tdT* into either mPOA or BNST to deplete Tph2 (Fig. 3a). As a control, we injected the same virus into the same regions of wild type (*WT*) mice. Co-staining of Tph2 and aromatic amino acid decarboxylase (AADC)––an enzyme required for biosynthesis of serotonin and dopamine––showed high overlap in control mice with Cre, but minimal overlap in *Tph2^flox/flox^* mice despite normal AADC expression (Fig. 3b, c). Moreover, terminal serotonin levels were significantly reduced at the injection site (Extended Data Fig. 4a), confirming the effectiveness and specificity of viral-mediated *Tph2* knockout. To systematically identify the function of 5HT_mPOA_ and 5HT_BNST_, we again tracked each female through different life stages, including pup-directed interactions both as a virgin and a mother (Fig. 3d).

**Fig. 3.**
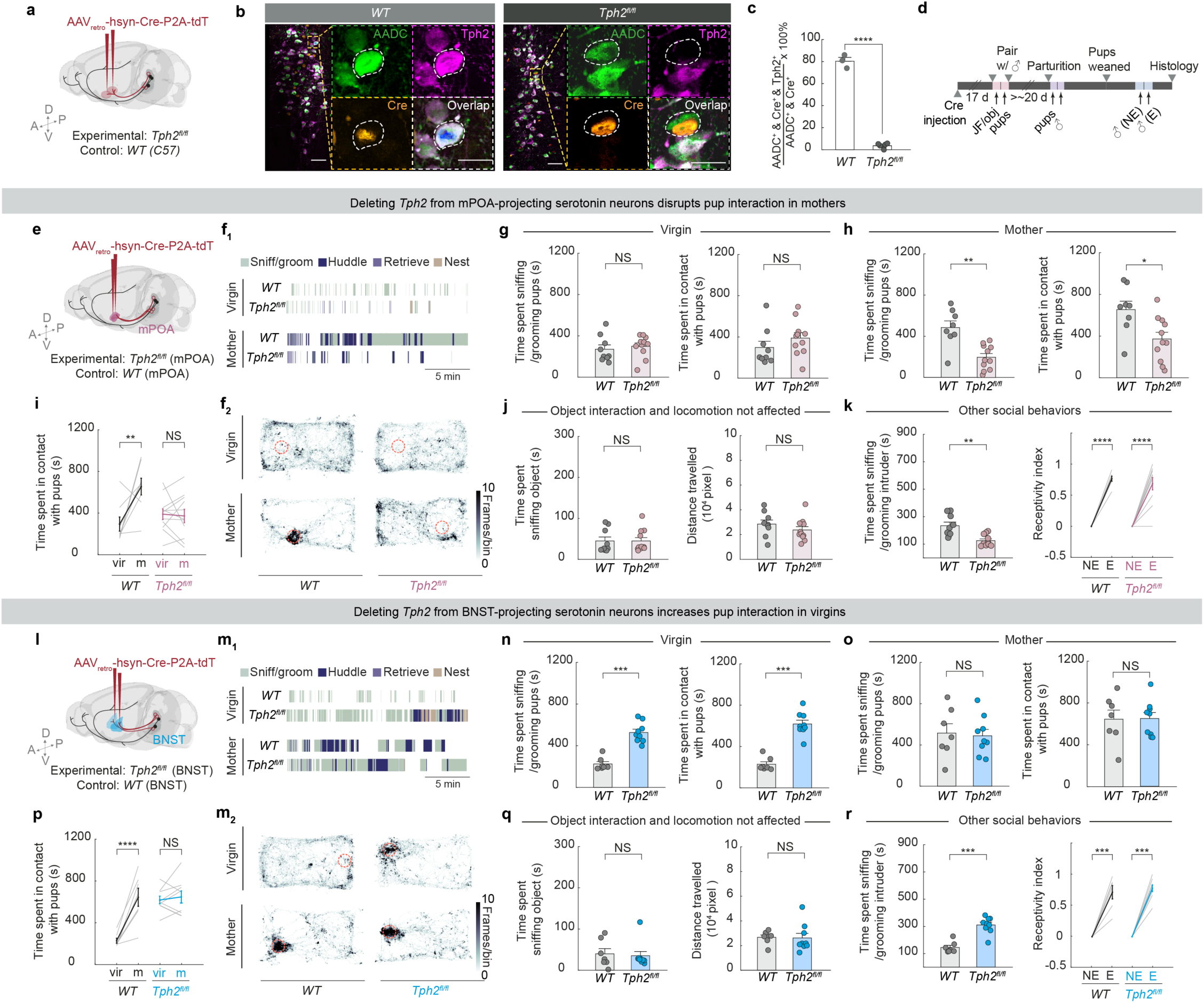
Serotonin is required in 5HT_mPOA_ to promote pup care in mothers, but in 5HT_BNST_ to inhibit pup interaction in virgins. **a**, Schematic showing viral-genetic strategy to conditionally knock out *Tph2* from projection-defined serotonin neurons. **b**, Representative histology showing that Tph2 (magenta) is expressed in AADC^+^ (green) neurons expressing Cre (orange) in wild type (*WT*) control mice (left), but is absent in *Tph2^flox/flox^*mice (right). Scale bars, 50 μm (20 μm in enlarged view of neurons outlined in the boxed areas). **c**, Percentage of AADC^+^ and Cre^+^ neurons that co-express Tph2 in *WT* and *Tph2^fl/fl^* mice (*n* = 3, 6; two-tailed *t-*test). The ∼20% AADC^+^ / Tph2^−^ cells in *WT* likely represent dopamine neurons near the DR. **d**, Experimental timeline. JF, juvenile female; obj, object; NE, non-estrus; E, estrus. Arrows indicate behavioral recording days (d). **e**, Schematic of control and 5HT_mPOA_-cKO neurons via bilateral injection of *AAV_retro_-Cre* into mPOA. **f**, Example ethogram showing pup-directed behaviors by a representative control and 5HT_mPOA_-cKO female as virgin (**f_1_**, top) vs. mother (**f_1_**, bottom). Example heat map showing the nose position of the same animals during pup interaction as virgin (**f_2_**, top) vs. mother (**f_2_**, bottom). Nose position was obtained using automatic animal pose tracking. Red circles indicate the main pup location in that session. **g**, **h**, Time spent sniffing/grooming pups (**g**, left; two-tailed *t-*test) or in contact with pups (**g**, right; Mann Whitney U*-*test) by mPOA-injected *WT* control and 5HT_mPOA_-cKO as virgins (*n* = 9, 11), and as mothers (**h**; two-tailed *t-* test) (8, 11; one control animal died during labor). **i**, Total pup contact time across reproductive stages in control and 5HT_mPOA_-cKO females (mixed effects analysis followed by Šídák’s multiple comparisons test). **j**, Novel object interaction (left) and distance travelled (right) by control and 5HT_mPOA_-cKO (*n* = 9, 11; object: Mann Whitney U-test; distance: two-tailed *t-*test). Distance travelled was calculated based on nose position obtained using automatic animal pose tracking. **k**, Left, time spent interacting with juvenile female intruder (*n* = 9, 11; two-tailed *t-*test) by mPOA-injected *WT* control and 5HT_mPOA_-cKO females. Right, receptivity across estrus cycle (right; *n* = 7, 10; mixed effects analysis followed by Šídák’s multiple comparisons test). **l–r**, Same as **e–k** except for 5HT_BNST_-cKO (*n* = 7, 9). For **c**, **g–k**, and **n–r**, each dot or line represents one animal; data are shown as mean ± s.e.m. See Source Data for detailed statistics.

In females with *Tph2* conditionally knocked out from 5HT_mPOA_ (5HT_mPOA_-cKO mice), the amount of pup interaction was comparable to controls as virgins (Fig. 3e–g; Extended Data Fig. 4i). However, as mothers, 5HT_mPOA_-cKO mice spent significantly less time sniffing, grooming, and maintaining physical contact with pups than controls despite normal pup retrieval (Fig. 3f, h; Extended Data Fig. 4c–e, j; Supplementary Video 3). The severity of the phenotype correlated with the number of cells with *Tph2* conditionally knocked out (Extended Data Fig. 4b, left). Pups raised by these 5HT_mPOA_-cKO mice were also smaller at weaning age compared to those raised by control mice despite similar litter size (Extended Data Fig 4k–n), suggesting that the deficit in maternal care was persistent. As control mice transitioned from virgins to mothers, engagement in pup interaction increased significantly (Fig. 3i, Supplementary Video 2). However, in 5HT_mPOA_-cKO mice, this state-dependent behavioral switch was abolished, as pup interaction remained low across reproductive states (Fig. 3i). Thus, serotonin is required in 5HT_mPOA_ to promote pup care in mothers.

### 5HTBNST suppress pup care in virgins

In striking contrast to Tph2 depletion from 5HT_mPOA_, Tph2 depletion from 5HT_BNST_ (5HT_BNST_-cKO) caused females to display heightened pup interaction in virgins (Fig. 3l–n; Extended Data Fig. 4f, i). Compared to controls, they spent significantly more time sniffing and grooming pups and maintaining physical contact with pups. Interestingly, however, 5HT_BNST_-cKO mice did not retrieve pups more than controls. Rather, they spent more time dragging nesting materials across the cage to re-nest around the pups (Extended Data Fig. 4g; Supplementary Video 4). Whereas the BNST-injected control mice showed a significant increase in pup interaction as mothers compared to themselves as virgins, 5HT_BNST_-cKO mice exhibited consistently high pup interaction regardless of reproductive states (Fig. 3o, p). Thus, serotonin is required in 5HT_BNST_ to suppress pup care in virgins.

To determine whether differences in parenting behavior could be attributed to more general behavioral deficits, we assessed each animal’s interactions with a novel object and baseline locomotion. Neither 5HT_mPOA_-cKO nor 5HT_BNST_-cKO mice differed significantly from control groups (Fig. 3j, q), indicating that the behavioral differences we observed in pup interaction were likely specific to social dynamics, rather than a secondary effect of altered locomotion or object interactions. Notably, 5HT_mPOA_-cKO mice interacted less with juvenile female intruders, whereas 5HT_BNST_-cKO mice interacted more than their respective controls, even though the level of social engagement initiated by the intruders did not differ significantly across experimental and control groups (Fig. 3k, r, left; Extended Data Fig. 4o, p). Yet, in the context of sexual behaviors, both 5HT_mPOA_-cKO and 5HT_BNST_-cKO mice showed normal state-dependent behavioral receptivity across estrus cycles (Fig. 3k, r, right). Therefore, the social phenotypes we observed are likely not specific to interactions with pups, but nonetheless do not generalize to all social behaviors.

Together, these data reveal profound functional heterogeneity within the serotonin system in regulating pup-directed behaviors. They demonstrate that collective actions of two distinct serotonin neuron populations— 5HT_mPOA_ and 5HT_BNST_ (Fig. 3i, p)—are required to coordinate the state-dependent switch in female pup care behavior.

### Sufficiency of 5HT_mPOA_ or 5HT_BNST_ in pup care or neglect

We next asked whether acutely activating 5HT_mPOA_ or 5HT_BNST_ is sufficient to change how females interact with pups. To selectively activate 5HT_mPOA_ or 5HT_BNST_, we injected an AAV_retro_ expressing Flp-dependent Cre into either mPOA or BNST of *Sert-Flp* mice, and a second AAV in the raphe nuclei (DR and MR) that expresses a step-function opsin with ultra-high light sensitivity (SOUL)^42^ in a Cre-dependent manner (Fig. 4a; Experimental). As one control, we injected the same viruses into *WT* mice (Fig. 4a; Control #1). During behavior tests, we introduced pups into a female’s home cage, allowed her to habituate for five minutes to attenuate novelty-driven exploration during stimulation, and then alternated every five minutes between blocks initiated by blue light stimulation, which activates SOUL (A), and orange light stimulation, which inactivates SOUL (I) (Fig. 4c; Extended Data Fig. 5a, b). The first block (blue or orange stimulation) was randomized across sessions. We tracked each female from virgins to mothers and assayed the effects of activating different projections on pup-directed behaviors (Fig. 4c).

**Fig. 4.**
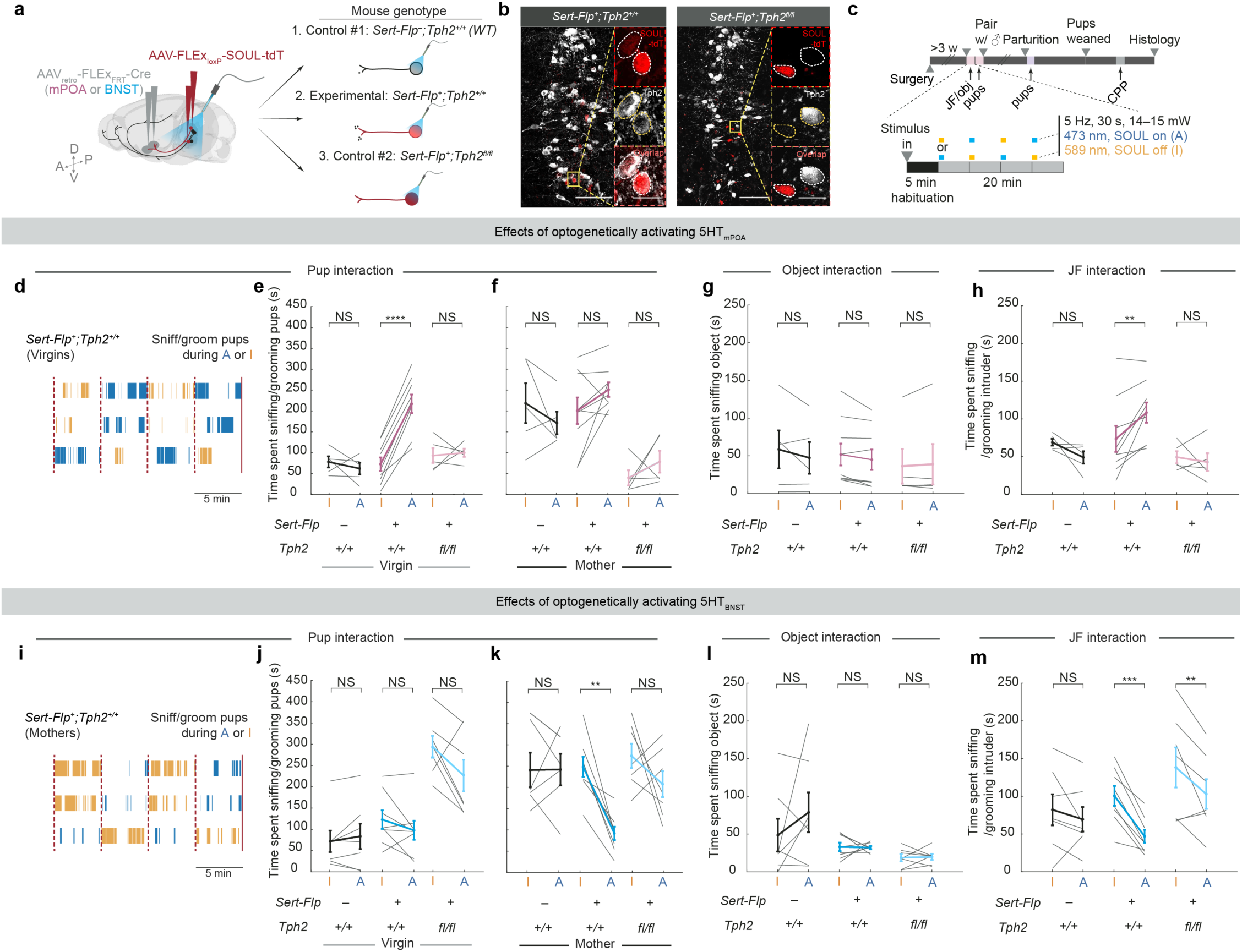
5HT_mPOA_ activation increases pup interaction in virgins, whereas 5HT_BNST_ activation inhibits pup interaction in mothers. **a**, Schematic showing viral-genetic strategy for (1) control #1, which have normal Tph2 expression but minimal SOUL expression as the animals lack Flp in Sert^+^ neurons; (2) experimental mice, which have SOUL (red) expressed in 5HT_mPOA_ or 5HT_BNST_ neurons with normal Tph2 expression (gray); and (3) control #2 with SOUL expressed in 5HT_mPOA_ or 5HT_BNST_ neurons that have *Tph2* conditionally knocked out. The three genotypes are presented in the same order from left to right in **e–h** and **j–m**. **b**, Representative confocal sections showing that SOUL expression (red) overlaps with Tph2 (gray) in *Sert- Flp^+^;Tph2^+/+^* mice (left, corresponding to Experimental in **a**), but does not in *Sert-Flp^+^;Tph2^fl/fl^*mice (right, corresponding to Control #2 in **a**). Scale bars, 100 μm (20 μm in high magnification inset). **c**, Experimental timeline and optogenetic stimulation paradigm for each behavioral session. CPP, conditioned place preference. Other abbreviations same as **Fig. 3d**. **d**, Three example ethograms of time spent sniffing/grooming pups during 5HT_mPOA_ activation in *Sert-Flp* virgins with normal Tph2. Each row is one mouse, with both SOUL off (I for inactive, orange) and SOUL on (A, for active, blue) periods shown. Red dotted lines on the ethograms mark the start of laser stimulations. Red solid lines mark the end of the session. **e**, **f**, Time spent sniffing/grooming pups during SOUL off (I) and SOUL on (A) by the mPOA-injected naïve virgin females (**e**) and as mothers (**f**). **g,** Object interaction between SOUL off (I) and SOUL on (A) in the mPOA-injected mice. **h**, Interaction time with a juvenile female intruder between SOUL off (I) and SOUL on (A) in the mPOA-injected mice. **i–m**, Same as **d–h** except for BNST-injected female mice. For **d–h** (*n* = 5, 9, 5) and **i–m** (*n* = 7, 8, 7 with one *WT* female that died during labor). Each dot or line represents one animal; data are shown as mean ± s.e.m. See Source Data for detailed statistics.

When we stimulated either the mPOA- or BNST-injected *WT* mice (Control #1), no significant difference was observed between orange and blue stimulation. As virgins, they exhibited low levels of pup interaction (Fig. 4e, j, left); as mothers, they showed significantly elevated pup interaction regardless of the stimulation (Fig. 4f, k, left). However, in experimental mice that express SOUL in 5HT_mPOA_ neurons, blue light stimulation significantly increased the amount of pup sniffing and grooming in naïve virgin females to a level that is comparable to *WT* mothers (Fig. 4d, e, middle; Extended Data Fig. 5c, e_1_). Yet, in lactating mothers, stimulating 5HT_mPOA_ no longer increased pup interaction significantly (Fig. 4f, middle), likely due to a ceiling effect caused by the vigorous care behavior endogenously displayed by mothers.

By contrast, stimulating 5HT_BNST_ had no effect on pup interaction in virgin females (Fig. 4j, middle), but significantly decreased pup interaction in mothers to a level comparable to that in virgins (Fig. 4i, k, middle; Extended Data Fig. 5d, e_2_). No pup-directed aggression was observed, suggesting that 5HT_BNST_ drive pup neglect but not infanticide.

Additionally, activating 5HT_mPOA_ neurons increased whereas activating of 5HT_BNST_ neurons decreased general social interaction with juvenile female intruders (Fig. 4h, m). Nevertheless, stimulation of neither experimental group induced significant changes in their interactions with a novel object (Fig. 4g, l), suggesting that the observed gain-of-function phenotypes are not a generic engagement effect, but likely a social effect.

### Gain-of-function effects require serotonin

Because the gain-of-function (GOF) experiment using *Sert-Flp* mice allows SOUL to activate the entire serotonin neuron, it is possible that other co-released neurotransmitters, instead of serotonin itself, drove the behavioral phenotypes we observed. To test the functional effects of activating the same neurons in the absence of serotonin, we performed a second control experiment—combining our GOF with a previous loss-of-function (LOF) approach—by injecting the same retrograde Flp-dependent Cre virus and Cre-dependent SOUL virus into *Sert- Flp;Tph2^flox/flox^* mice (Fig. 4a; Control #2). Under this paradigm, 5HT_mPOA_ and 5HT_BNST_ still express SOUL but their *Tph2* is conditionally deleted, preventing serotonin synthesis and release when the neuron is activated. Staining of Tph2 protein verified that Tph2 was indeed absent from SOUL^+^ cells (Fig. 4b).

Notably, mPOA-injected *Sert-Flp;Tph2^flox/flox^* mice (an independent way of producing 5HT_mPOA_-cKO from our previous manipulation) recapitulated the LOF phenotype observed in the previous 5HT_mPOA_-cKO mice (Fig. 3i): as mothers, these mice showed significantly lower levels of pup sniffing and grooming compared to *Sert-Flp* and *WT* mothers. Stimulating 5HT_mPOA_ lacking Tph2 could not rescue this LOF deficit (Fig. 4f, right). Moreover, unlike *WT* virgins, activating 5HT_mPOA_ lacking Tph2 failed to increase pup interaction in virgin females (Fig. 4e, right). This provided strong evidence that serotonin itself is the principal player in these 5HT_mPOA_ neurons to promote pup interaction.

Conversely, BNST-injected *Sert-Flp;Tph2^flox/flox^* mice spent significantly more time sniffing and grooming pups as virgins at levels comparable to *Sert-Flp* and *WT* mothers, replicating the phenotype observed in previous 5HT_BNST_-cKO mice (Fig. 4j, right; compared to Fig. 3p). Yet, although there was a trend of reduced pup interaction by SOUL stimulation, the heightened pup interaction in *Sert-Flp;Tph2^flox/flox^* virgins could not be fully rescued by activating 5HT_BNST_ lacking Tph2 (Fig. 4j, right). Further, in contrast to the GOF alone in *Sert-Flp* mice, which suppressed pup interaction in mothers, activation of 5HT_BNST_ lacking Tph2 no longer suppressed pup care (Fig. 4k, right), indicating that the GOF effect depends on serotonin. Nevertheless, the same activation did significantly reduce general social interaction with juvenile females (Fig. 4m, right), though not to a level as low as activation in the *Sert-Flp* condition. Thus, although additional neurotransmitters may contribute, serotonin is a key contributor in the 5HT_BNST_ neurons in suppressing social interactions.

### 5HTmPOA activation is rewarding

Serotonin has been implicated in both reward and punishment^18,21,24,38,43,44^. To examine whether valence contributed to the changes of pup interaction induced by loss- and gain-of-function manipulations, we performed conditioned place preference tests on control (Control #1), SOUL expression alone (Experimental), or SOUL expression in cKO background (Control #2) for both 5HT_mPOA_ and 5HT_BNST_ (Extended Data Fig. 5g). None of the groups avoided the chamber with blue light stimulation (Extended Data Fig. 5h, i), suggesting that activation of 5HT_mPOA_ or 5HT_BNST_ were not aversive. Hence, suppression of pup interaction induced by 5HT_BNST_ activation is unlikely a byproduct of animals finding the activation aversive.

Interestingly, mice with SOUL expressed in 5HT_mPOA_ showed a significant increase in preference for the chamber with blue light stimulation, and this preference was abolished in the cKO background (Extended Data Fig. 5h, i), suggesting that serotonin release from 5HT_mPOA_ neurons is rewarding. Given the importance of care behavior, it is possible that this coupling of care promotion and reward in 5HT_mPOA_ is evolutionarily selected such that caring for pups is innately rewarding in mothers.

### Dynamic 5-HT release across states

We next consider whether our fiber photometry recording of Ca^2+^ signals from the soma of serotonin neurons (Fig. 1) and loss- and gain-of-function results (Fig. 3 and Fig. 4) could be accounted for by changes in serotonin release across a female’s reproductive stages. To test this hypothesis, we injected serotonin sensor (g5-HT3.0)^45^ into either mPOA or BNST and recorded serotonin dynamics using fiber photometry as the females interacted with pups, object or a juvenile female intruder before and after parturition (Fig. 5a, g).

**Fig. 5.**
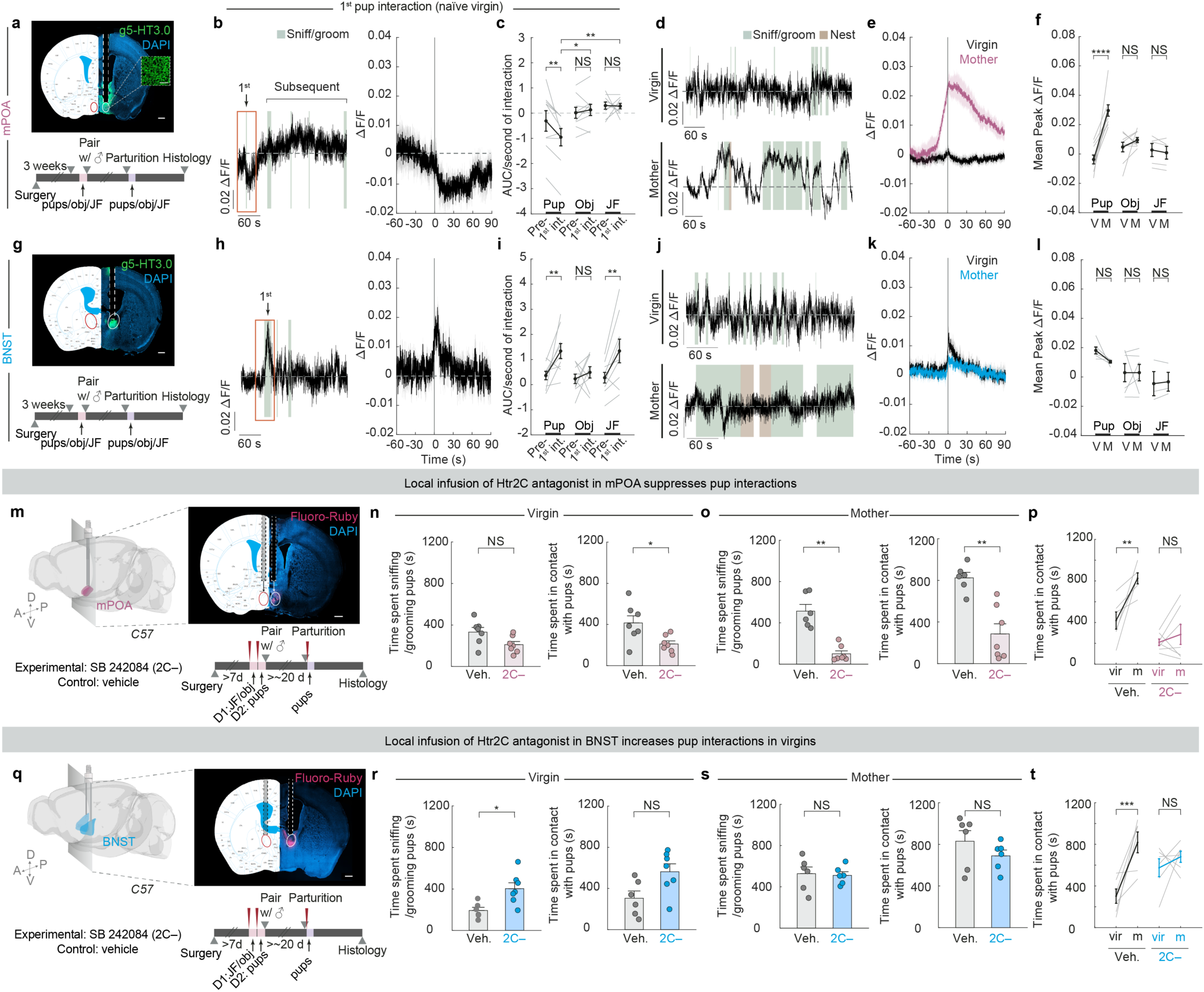
Local dynamics and action of serotonin at mPOA and BNST. **a**, Top, representative histology aligned to the mouse brain atlas^54^ showing fiber implantation for serotonin sensor (g5-HT3.0) recording in mPOA. Bottom, experiment timeline. All animals were recorded at two wavelengths: 490 nm which captures serotonin-dependent fluorescence and 405 nm, an isosbestic control that captures serotonin-independent background noise and motion artifacts. Scale bars, 500 μm (50 μm in high magnification inset) **b**, Left, representative Δ*F*/*F* trace of g5-HT3.0 signal in mPOA during the first and subsequent pup interactions in a naïve virgin. Right, group g5-HT3.0 signal in mPOA during the first pup interaction in naïve virgins. *n* = 9 recorded females. **c**, Quantifications of area under curve (AUC) normalized to interaction duration before and during the first pup, object, and juvenile female interaction in naïve virgins. (*n* = 9 females recorded in mPOA; two-way repeated measures (RM) ANOVA followed by Tukey multiple comparisons tests). **d**, Representative Δ*F*/*F* traces of g5-HT3.0 signal in mPOA during the pup interactions in a naïve virgin (top) and the same animal after parturition (bottom). **e**, Peri-event time histograms (PETHs) of Δ*F*/*F* of g5-HT3.0 signal in mPOA aligned to the onset of females sniffing/grooming pups. The panel shows both group g5-HT3.0 signal in naïve virgins (black trace) and the same females as mothers (pink trace) (*n* = 9 females). **f**, Within-animal comparisons of mean peak Δ*F*/*F* of g5-HT3.0 signal in mPOA between virgins (V) and the same animals as mothers (M) during pup, object, and juvenile female (JF) interaction (*n* = 9 virgins and mothers except *n* = 5 for JF interaction; the other 4 mothers attacked the juvenile female and were excluded as the behavior was not comparable to their non-aggressive interactions with juvenile female intruder as virgins; two-way RM ANOVA followed by Tukey multiple comparisons tests). **g–l,** Same as **a–f**, but for g5-HT3.0 recording at BNST. *n* = 9 (**h, i**); *n* = 6 (**k, l**) except *n* = 3 for JF interaction in (**l**). **m**, Experimental setup for Htr2C antagonist (2C–) infusion in mPOA and representative histology showing infusion target indicated by Fluoro-Ruby (magenta). Scale bar, 500 μm. **n**, Time spent sniffing/grooming pups (left; *n* = 7, 7; two-tailed *t-*test) or maintaining contact with pups (right; *n* = 7, 7; two-tailed *t-*test) by naïve virgin females with 2C– or vehicle infused in mPOA. **o**, Time spent sniffing/grooming pups (left; *n* = 6, 7 Mann Whitney U test) or maintaining contact with pups (right; *n* = 6, 7, one control female died during labor; two-tailed *t-*test) by mothers with 2C– or vehicle infused in mPOA. **p,** Total pup contact time across reproductive stages in mice with 2C– or vehicle infused in mPOA (*n* = 7, 7 for virgins; *n* = 6, 7 for mothers; mixed effects analysis followed by Šídák’s multiple comparisons test). **q–t,** Same as **m–p** except for 2C– infusion in BNST. *n* = 6, 7 for vehicle and 2C– (**r**); *n* = 6, 6 for vehicle and 2C– (one 2C– female died during labor); *n* = 6, 7 for virgins and *n* = 6, 6 for mothers (**s, t**). See Source Data for detailed statistics.

Surprisingly, upon the first interaction with pups as naïve virgins, serotonin signal recorded in mPOA dropped significantly (Fig. 5b). This decrease was seen across all nine animals recorded and was specific to the first pup interaction, as it was not observed during the first interaction with an object or a juvenile female, or subsequent pup interactions (Fig. 5b–d; Extended Data Fig. 6a, left). By contrast, serotonin signal in BNST rose sharply upon the first pup interaction as well as subsequent interactions (Fig. 5h–j; Extended Data Fig. 6a, right). During the first juvenile female interaction, serotonin signal in BNST also increased significantly (Fig. 5i), an observation consistent with previous single-cell Ca^2+^ imaging of BNST-projecting serotonin neurons in DR^25^. No significant increase was detected in either the first or subsequent object interactions (Fig. 5i; Extended Data Fig. 6d). These results indicate that the serotonin release in BNST is likely a social signal.

Although there was no substantial change in serotonin signal in mPOA when naïve virgins interacted with pups after the first encounter, we observed recurring and sustained bouts of serotonin release in the same females as lactating mothers, which initiated before but peaked at the onset of pup interaction on average (Fig. 5d, e). The magnitude of change in serotonin activity in mPOA in mothers was significantly higher than when the same animals were virgins (Fig. 5e, f). Such change in sensor signal was unlikely due solely to difference in viral expression or motion artifacts, as serotonin release in response to an object or a juvenile female did not differ significantly from virgins to mothers, nor did we observe the same pup-directed signal increase when we recorded using the mutant version of g5-HT3.0 in mPOA (Fig. 5f; Extended Data Fig. 6b, c, f–i). In the BNST, the peak ΔF/F of serotonin signal in response to pups did not change significantly across reproductive stages (Fig. 5k, l; Extended Data Fig. j–m).

Together, these observations revealed opposing serotonin release patterns at the sites of mPOA and BNST in response to pups during the first pup interaction in naïve virgins. Furthermore, changes in serotonin release at mPOA––from a significant decrease upon the first ever pup encounter in virgins to a sustained, high level serotonin response towards the same type of social stimulus in mothers––accentuate mPOA as a cardinal, dynamic locus of serotonin release with considerable basal level of serotonin activity. Such dynamics make serotonin in mPOA poised to bidirectionally tune pup-directed care behavior.

### Local action of 5-HT at mPOA and BNST

Individual serotonin neurons often collateralize to multiple brain regions^18,19,46,47^. Therefore, our loss- and gain-of-function experiments thus far could not rule out the possibility that the action of 5HT_mPOA_ or 5HT_BNST_ in modulating pup interaction is at brain regions that receive collateralized projections from these neurons, rather than at mPOA or BNST themselves. To investigate whether serotonin activity is required locally in mPOA or BNST to regulate state-dependent pup care, we took a pharmacological approach to locally infuse serotonin receptor antagonists to interfere with serotonin signaling. As serotonin receptor 2C (Htr2C) is consistently one of the most highly expressed serotonin receptors at the transcriptomic level in both regions based on previous single-nucleus^48^ and single-cell sequencing studies^49,50^, we infused SB 242084^51,52^, an Htr2C antagonist (hereafter, 2C–), directly into mPOA or BNST. As a control, we infused the same volume of vehicle into the same brain regions of the littermates (Fig. 5m, q).

When 2C– was infused into the mPOA, females interacted significantly less with pups as both virgins and mothers (Fig. 5n, o). Whereas the vehicle-infused mice showed a significant increase in pup interaction from virgins to mothers, females with 2C– infused in mPOA displayed consistently low levels of pup interaction (Fig. 5p). Moreover, these 2C– infused mothers showed significant deficits in pup retrieval (Extended Data Fig. 7b, right). By contrast, when the same 2C– was infused into BNST, virgin females spent significantly more time sniffing and grooming pups at levels comparable to mothers (Fig. 5r–t). No significant effect was observed when 2C– was infused into BNST in mothers (Fig. 5s; Extended Data Fig. 7f). As Htr2C is an excitatory serotonin receptor^53^, these results are consistent with serotonin promoting and suppressing pup interaction through excitatory effects on target neurons at mPOA and BNST.

The consistencies between the *Tph2* conditional knockout and 2C– infusion experiments (Extended Data Fig. 8) demonstrate that serotonin acts locally at mPOA to promote pup interaction in mothers and at BNST to inhibit pup interaction in virgins. Furthermore, the fact that acute Htr2C inhibition gave similar results as conditional knockout of *Tph2* weeks before the behavioral assays (Fig. 3d) suggests that the effects of conditional knockout were caused by acute loss of serotonin rather than compensatory changes due to long-term serotonin deprivation. Additionally, although other serotonin receptors could also contribute, these experiments suggest Htr2C as a key receptor that mediates serotonin signaling in both mPOA and BNST neurons to modulate pup care behavior.

## Discussion

The transition from a nonparent to a parent entails drastic physiological and behavioral changes^5^. Whereas naïve virgin females do not and do not have the need to care for offspring, mothers invest significant time and metabolic resources to ensure the survival of the young. This fundamental transition is a hallmark of efficient and adaptive animal behavior, and must be supported by underlying changes at the neural level. Here, we reveal a parallel, segregated circuit logic of serotonin modulation that effectuates this flexibility in females’ pup-directed behaviors across reproductive stages.

During pup interactions, multiple subsystems of serotonin neurons, defined by their projection targets, are recruited. Contrary to the traditional view that serotonin invariably promotes sociability, our data demonstrate clear functional heterogeneity of serotonin subsystems in pup-directed behaviors. At the circuit level, serotonin from 5HT_mPOA_ promotes pup interaction, whereas serotonin from 5HT_BNST_ suppresses it. At the molecular level, antagonizing the same excitatory serotonin receptor 2C at mPOA and BNST similarly led to opposite behavioral effects on pup-directed behaviors, consistent with the known functions of these two regions in promoting and suppressing pup care^4,34,36^. At the physiological level, serotonin activity has distinct dynamics within mPOA and BNST during pup interactions, indicating that serotonin subpopulations can exert independent control of downstream target regions. Equipped with such parallel anatomical access to functionally distinct nodes of the care circuit, serotonin from distinct subsystems is poised to up- or downregulate pup interactions.

Importantly, even though serotoninergic circuits responsible for suppressing (5HT_BNST_) and promoting (5HT_mPOA_) pup care are in place in both virgins and mothers, distinct behavioral responses towards pups can be generated from the same circuit architecture as serotonin release changes. In mPOA, serotonin activity shifts across reproductive stages, decreasing upon the first pup encounter to potentially reinforce 5HT_BNST_ in discouraging care in naïve virgins through the mPOA–BNST reciprocal inhibitory circuits^4^, but increasing in lactating mothers to promote and reinforce care. Although 5-HT activity in BNST did not mirror the change observed in mPOA, it is possible that serotonin alters the balance within the mPOA–BNST circuits^4^ by leveraging existing connectivity, where the elevated serotonin release in mPOA indirectly decreases BNST activity through increased inhibition from mPOA. Consequently, latent pup care behavior is actively suppressed in naïve virgins by serotonin release from 5HT_BNST_, but becomes unleashed when 5HT_mPOA_ has sufficiently upscaled its serotonin activity during motherhood. Together, these results highlight serotonin release as a mechanism of plasticity, accentuate the dynamic role of serotonin in tuning neural dynamics to toggle between different behavioral outputs, and reveal a circuit mechanism where joint actions of distinct serotonin neurons coordinate adaptive behavior changes across life stages.

---

**Extended Data Fig. 1.**
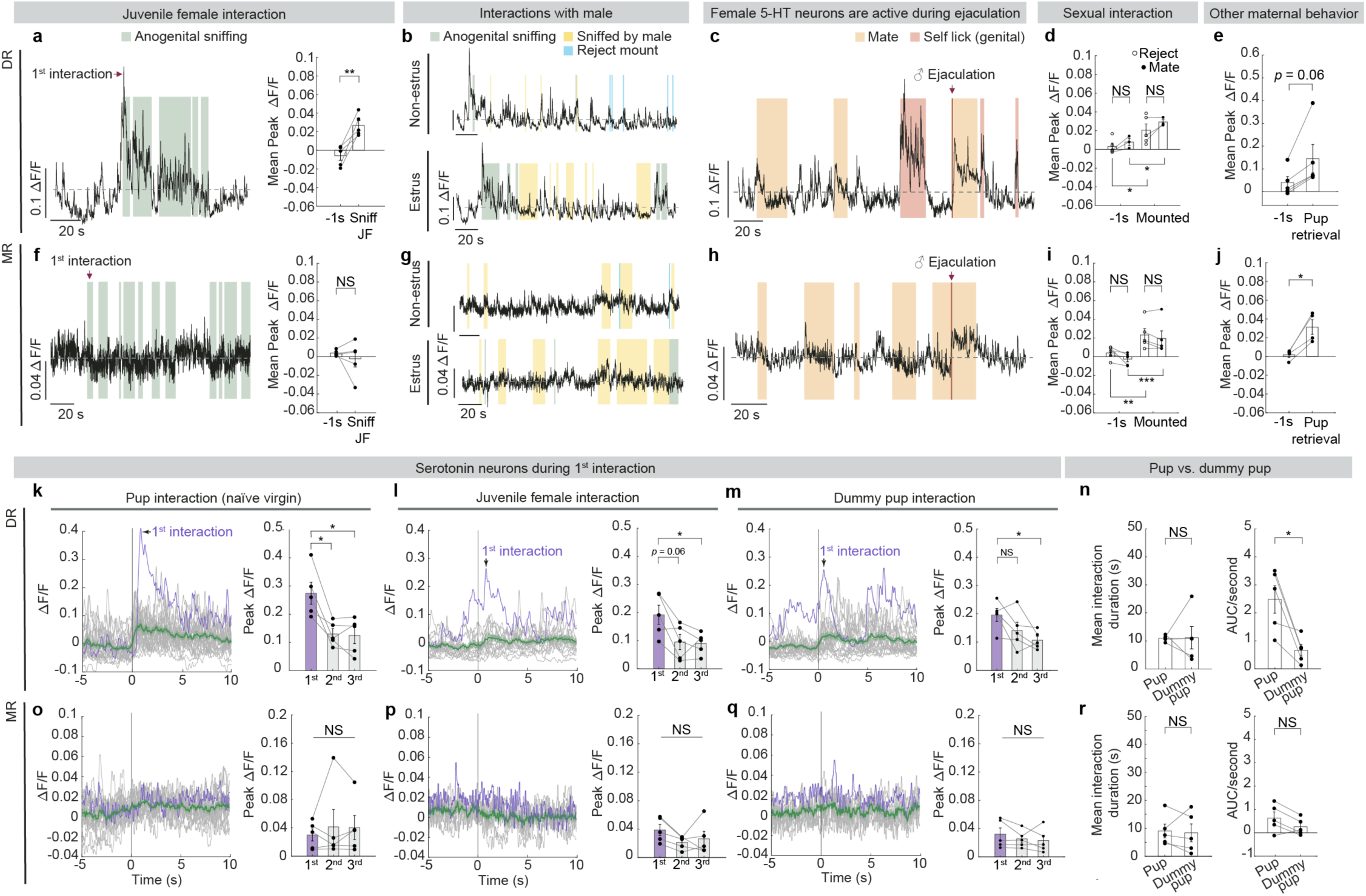
Additional characterization of serotonin neuron activity during social interactions. **a**, Left, representative Δ*F*/*F* traces of DR serotonin neurons during interactions with a juvenile female intruder; Gray dash lines in these and all subsequent panels indicate Δ*F*/*F* = 0. Right, quantification of mean peak Δ*F*/*F* before and during juvenile female interactions (*n* = 5 females; paired *t-*test). Female DR serotonin neurons showed significant increase in activity during interactions with a juvenile female. **b**, Top, representative Δ*F*/*F* traces of DR serotonin neurons during interactions with a male intruder when the recorded female was in non-estrus. Bottom, representative Δ*F*/*F* traces of DR serotonin neurons during interactions with a male intruder when the recorded female was in estrus. **c**, Representative Δ*F*/*F* traces of DR serotonin neurons in females during mating and when the male ejaculated. Serotonin neurons in female mice showed increase in activity following male ejaculation (single ejaculation trial recorded from *n* = 3 females in DR, and *n* = 2 females in MR, see panel **h**), consistent with a previous study reporting that female DR serotonin neurons are activated by ejaculation^26^. **d**, Quantification of mean peak Δ*F*/*F* of DR serotonin neurons before and during male mounting attempts. Hollow circles indicate when the female rejected mounting during non-estrus, and solid circles indicate when the female mated with the male during estrus (*n* = 5 females; note that 2 out of 5 DR females did not mate during estrus recordings, mixed effects analysis performed on the three females recorded during both estrus cycles followed by Šídák’s multiple comparisons test). No significant change in female serotonin neuron activity was observed at the population level during sexual interactions with male across estrus cycle. **e**, Quantification of mean peak Δ*F*/*F* before and during pup retrieval in mothers (*n* = 5 females; Wilcoxon matched-pairs signed rank test). **f–j**, Same as **a–e**, but in MR (*n* = 5 for all except *n* = 4 for panel **j**; one mother did not retrieve). **k**, Left, PETHs of Δ*F*/*F* of a representative naïve virgin female during pup interactions recorded in the DR. Right, group peak Δ*F*/*F* during the first three pup interactions (one-way ANOVA). The first pup interaction triggered significantly higher activity in DR serotonin neurons than following interactions. Purple trace, the first interaction. Green trace, mean response of all interactions ± s.e.m. **l**, **m**, Same as (**k**) but for juvenile female (**l**) or dummy pup (**m**) interactions. **n**, Left, mean interaction duration with real pups and dummy pups in DR-recorded naïve virgin females. Right, interaction-normalized activity of DR serotonin neurons during interactions with real pups or dummy pups in naïve virgin females (paired *t-*test). **o–r**, Same as **k–n**, but in MR (*n* = 5). See Source Data for detailed statistics.

**Extended Data Fig. 2.**
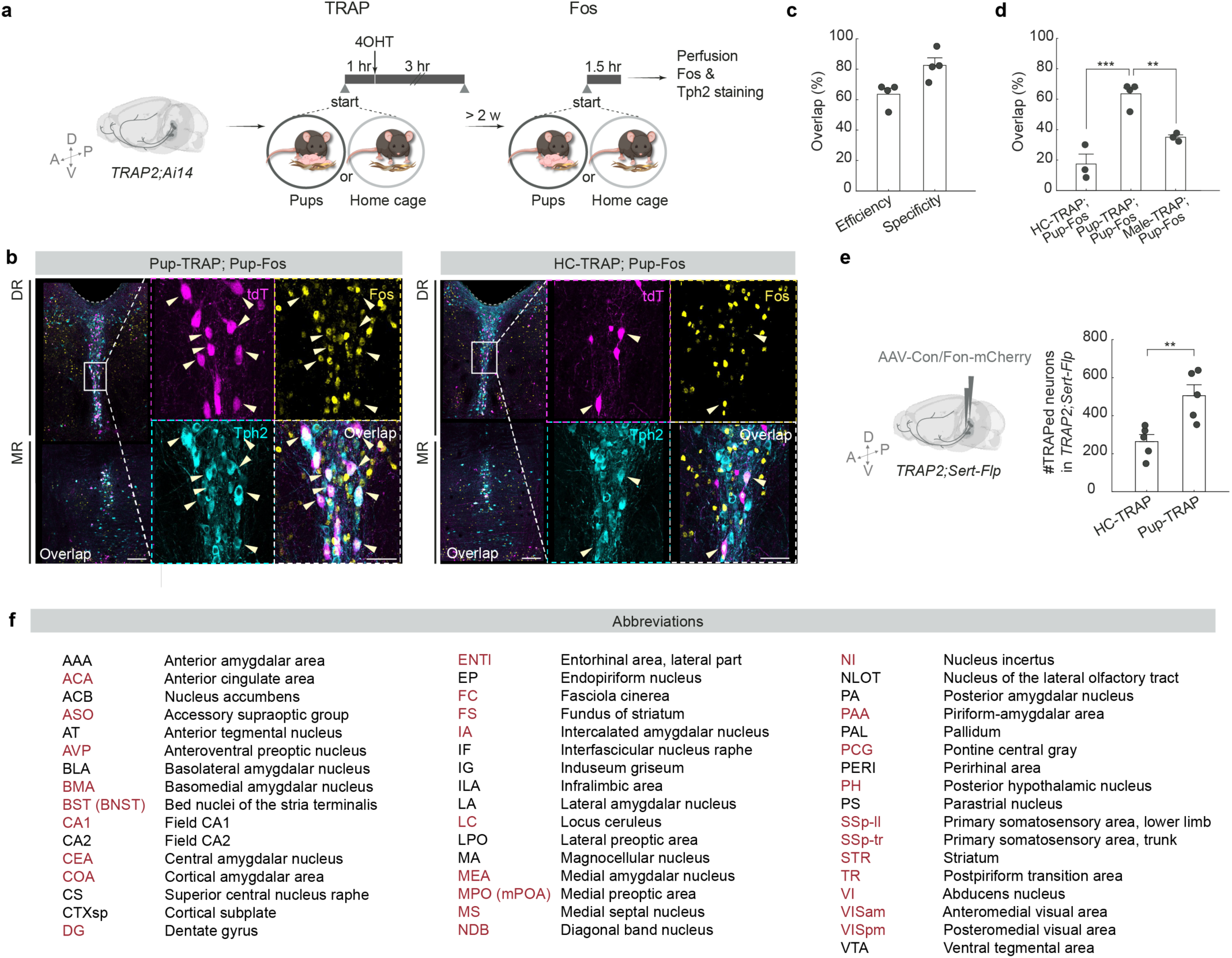
Validation of TRAP in capturing pup-activated serotonin neurons. **a**, Schematic of viral-genetic strategy for validating TRAP in capturing pup-activated serotonin neurons. **b**, Representative histology showing the overlap of TRAP-induced tdTomato expression (magenta), Fos (yellow) and Tph2 staining (cyan) from a naïve virgin female subjected to pup interaction both during TRAP and before Fos staining (left) and a naïve virgin female subjected to home cage interaction during TRAP and pup interaction before Fos staining (right). Arrows point towards example tdTomato^+^ cells that are Tph2^+^. Scale bars, 100 μm (50 μm in enlarged view of neurons outlined in the boxed areas). **c**, The efficiency and specificity of TRAP in capturing pup-activated serotonin neurons (*n* = 4). Efficiency = #triple^+^ / #Fos^+^ & Tph2^+^. Specificity = #triple^+^ / #tdTomato^+^ & Tph2^+^. These data indicate that the majority of pup-TRAPed neurons are also activated by pup interaction measured by Fos immunostaining, and vice versa. **d**, Percentage of TRAP/Fos overlap in Tph2^+^ neurons (*n* = 3 HC-TRAP and pup-Fos, 4 pup-TRAP and pup-Fos, 3 male-TRAP and pup-Fos; one-way ANOVA), calculated as #triple^+^ / #Fos^+^ & Tph2^+^. These data indicate that pup-TRAP can preferentially capture serotonin neurons that are activated by pup interactions. **e**, The number of labeled neurons is significantly higher in the pup-TRAP than the HC-TRAP conditions from Fig. 2 (*n* = 5, 5; two-tailed *t-*test). **f**, Allen abbreviations and full names of all brain regions that exhibit significantly higher raw axon density labeling in the pup-TRAP condition than the HC-TRAP condition, at 3% false discovery rate. Brain regions that are also among the top 30 regions that differ between the two conditions in normalized labeling density are highlighted in red. Regions are listed in alphabetical order. Note that in the current paper, Allen abbreviation BST is denoted as BNST, and MPO and MPN are referred to collectively as mPOA. See Source Data for detailed statistics.

**Extended Data Fig. 3.**
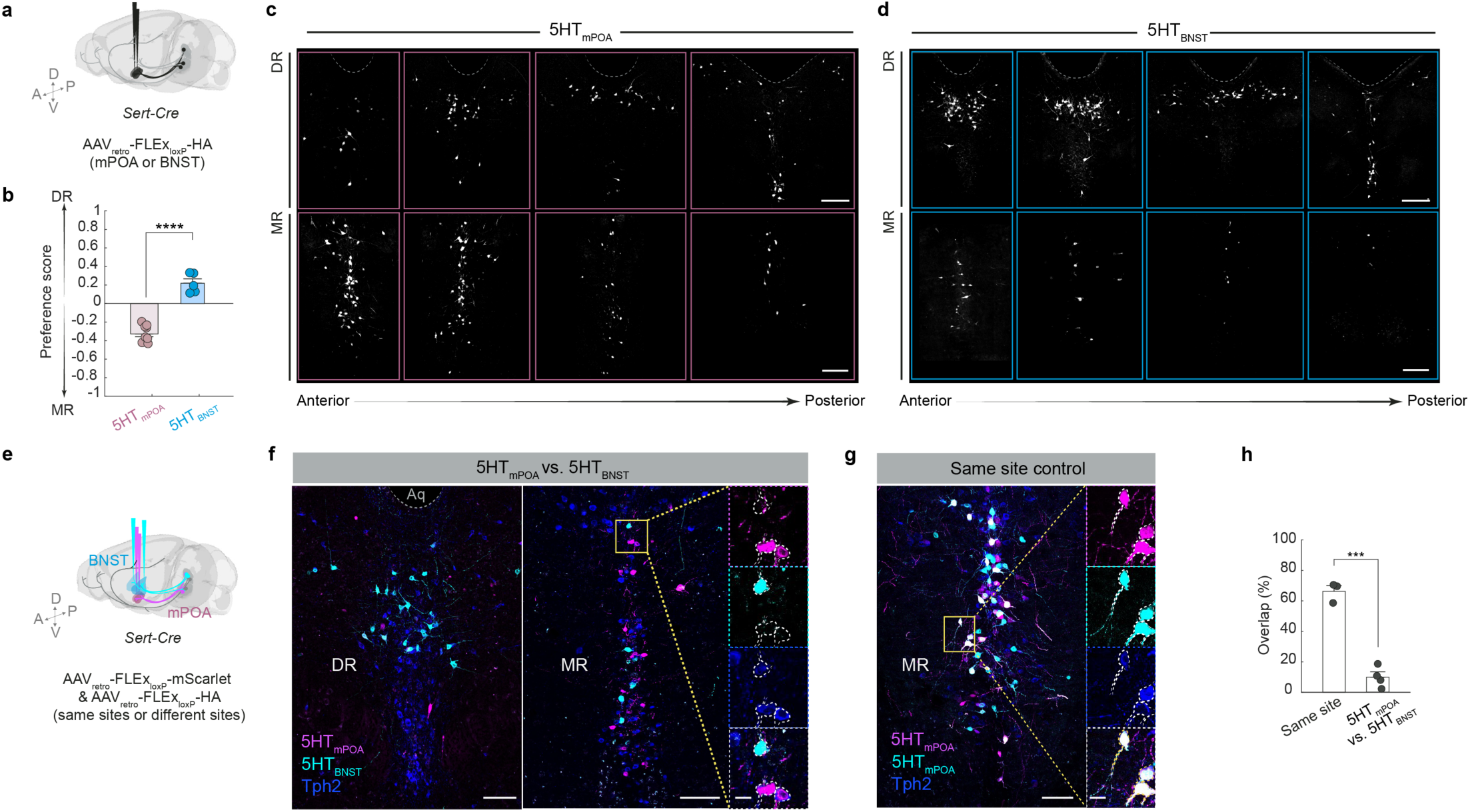
5HT_mPOA_ and 5HT_BNST_ show different cell body location preference in the raphe and little overlap. **a**, Schematic of viral-genetic strategy for retrogradely labeling 5HT_mPOA_ and 5HT_BNST_ neurons. **b**, Preference score of 5HT_mPOA_ and 5HT_BNST_ cell body location in the DR and MR (*n* = 7, 5; two-tailed *t-*test), calculated as (#DR – #MR) / (#DR + #MR). **c**, **d**, Representative histology showing the distribution of 5HT_mPOA_ (**c**) and 5HT_BNST_ (**d**) cell body location in the DR (top) and MR (bottom). Dotted gray lines outline the edge of the aqueduct. Scale bars, 200 μm. **e**, Schematic of viral-genetic strategy for dual labeling of 5HT_mPOA_ and 5HT_BNST_ neurons in the same brain. **f**, **g**, Representative histology showing the overlap of 5HT_mPOA_ (magenta) and 5HT_BNST_ (cyan) with Tph2 staining (blue) (**f**), and the overlap when both viruses were sequentially injected into mPOA as a control (**g**). Scale bars, 100 μm (20 μm in enlarged view of neurons outlined in the boxed areas). **h**, Percentage overlap between 5HT_mPOA_ and 5HT_BNST_ neurons is significantly lower than when the two retrograde Cre-dependent AAVs were injected into the same site (*n* = 3, 4; two-tailed *t-*test). See Source Data for detailed statistics.

**Extended Data Fig. 4.**
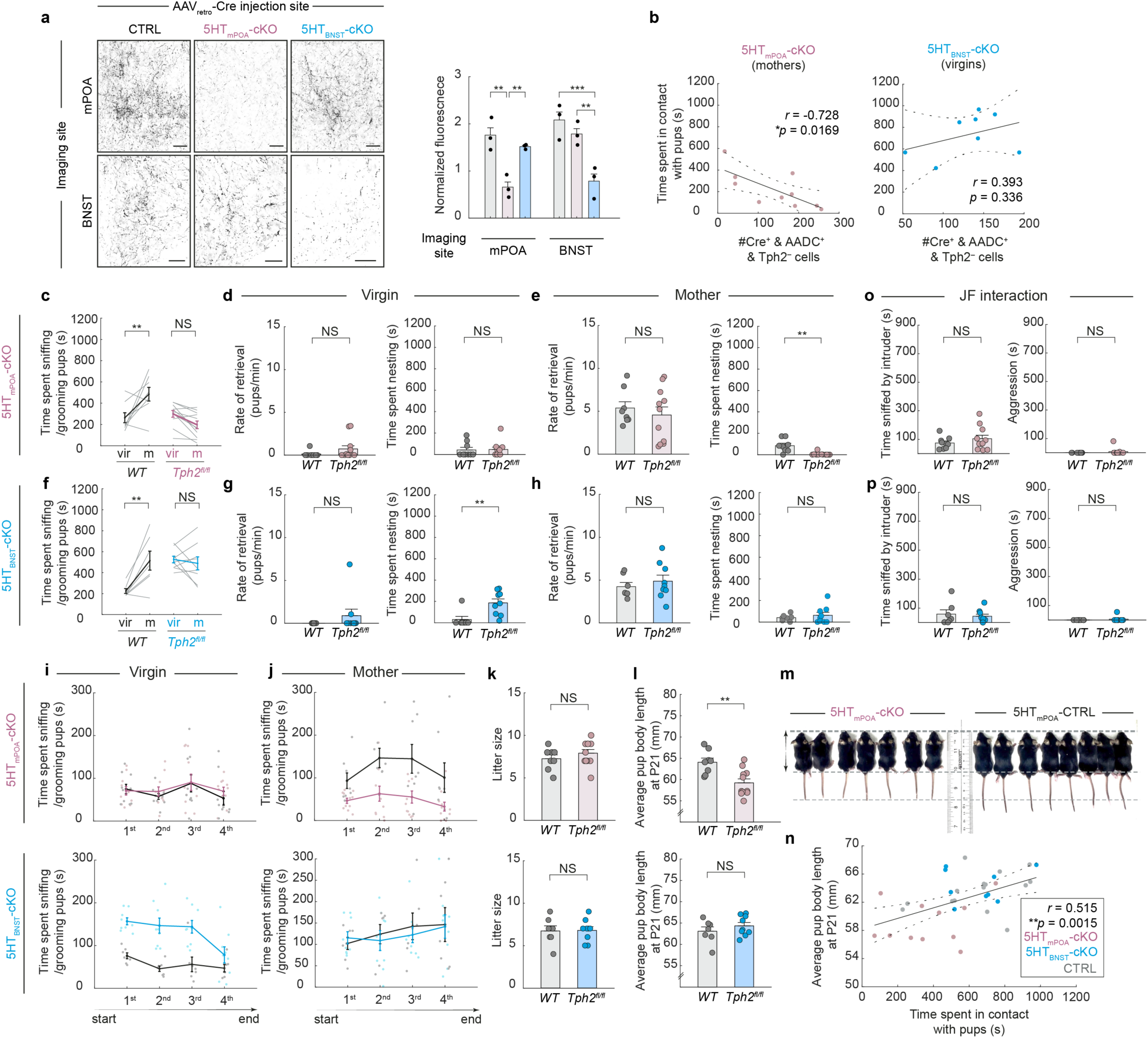
Additional characterization of 5HT_mPOA_-cKO and 5HT_BNST_-cKO histology and phenotypes. **a**, Representative serotonin staining in mPOA and BNST after *AAV_retro_-Cre* injection into either mPOA or BNST of *WT* or *Tph2^flox/flox^* mice (**a,** left). Normalized fluorescence of serotonin staining in mPOA and BNST in control (black), 5HT_mPOA_-cKO (pink) and 5HT_BNST_-cKO (blue) mice. Two-way ANOVA followed by Šídák’s multiple comparisons test (*n* = 3, 3, 3). Scale bars, 50 μm. **b**, Pearson correlation between the number of Cre^+^ and AADC^+^ but Tph2^−^ cells (that is, *Tph2* knockout serotonin neurons) and the total time spent in pup contact in 5HT_mPOA_-cKO mothers (left) or 5HT_BNST_-cKO naïve virgins (right). **c**, Time spent sniffing or grooming pups across reproductive stages in 5HT_mPOA_-cKO (*n* = 8, 11; one control animal died during labor; mixed effects analysis followed by Šídák’s multiple comparisons test). **d**, **e**, Rate of retrieval (left) and time spent nesting around pups (right) by control and 5HT_mPOA_-cKO mice as naïve virgins (**d**) and mothers (**e**). **f–h**, Same as **c–e** except for 5HT_BNST_-cKO. **i**, **j**, Time spent sniffing/grooming pups by 5HT_mPOA_-cKO females (top) and 5HT_BNST_-cKO females (bottom) as virgins (**i**) and mothers (**j**). 1^st^ through 4^th^ denote the first through last five-minute-long quarters within the single behavioral session. Pup interaction by 5HT_mPOA_-cKO mothers was consistently low throughout the behavioral session (**j**), whereas that by 5HT_BNST_-cKO virgins was high since the start of the session and remained so during most of the session (**i**). **k**, Litter size did not differ significantly between 5HT_mPOA_-cKO (top) or 5HT_BNST_-cKO females (bottom) and their respective controls. **l**, The average body size (from snout to tail base, indicated the arrows) at weaning age (postnatal day 21, P21) is significantly lower among pups raised by 5HT_mPOA_-cKO mothers (top), but not those raised by 5HT_BNST_-cKO mothers (bottom). **m**, Two example litters weaned from a 5HT_mPOA_-cKO mother (left) and an mPOA-injected control mother (right). **n**, The average body size of pups at wean date shows a significant positive correlation with the time spent in contact with pups in the recorded behavior sessions (control, gray; 5HT_mPOA_-cKO, pink; 5HT_BNST_-cKO, blue). **o, p,** The amount of social sniffing initiated by the juvenile female intruder (left) and the amount of aggression towards the juvenile female intruder do not differ significantly between 5HT_mPOA_-cKO females and their controls (**o**) or between 5HT_BNST_-cKO females and their controls (**p**). See Source Data for detailed statistics.

**Extended Data Fig. 5.**
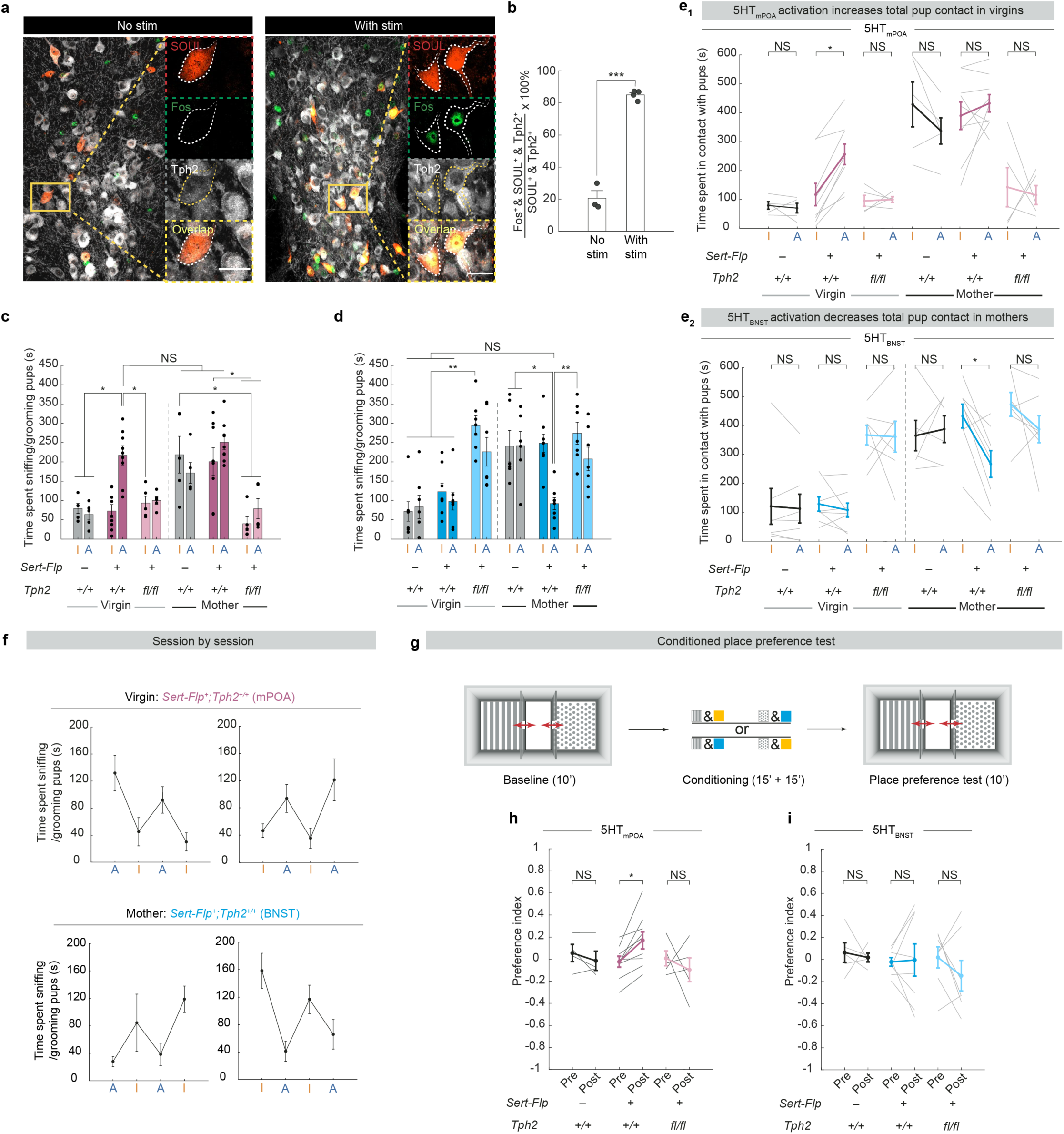
Validation of SOUL activation and additional behavioral characterizations. **a**, Representative histology showing the overlap of SOUL (red), Fos (green) and Tph2 (gray) without (left) and with (right) blue light stimulation. Scale bar, 20 μm. **b**, Percentage of Fos^+^ cells out of all SOUL^+^ serotonin neurons is significantly higher following blue light stimulation (*n* = 3, 4; two-tailed *t* test). **c**, **d**, Time spent sniffing/grooming pups during SOUL off (I) and SOUL on (A) by the mPOA-injected (**c**) and BNST-injected (**d**) mice. Statistics shown are from the same analyses as **Fig. 4e–f**, **j–k**. **e**, Time spent in contact with pups during SOUL off (I) and SOUL on (A) by the mPOA-injected (**e_1_**) and BNST- injected (**e_2_**) mice [mPOA: *n* = 5 *WT*, 9 *Sert-Flp^+^;Tph2^+/+^*, 5 *Sert-Flp^+^;Tph2^fl/fl^*; BNST: *n* = 7 *WT* (one died during labor), 8 *Sert-Flp^+^;Tph2^+/+^*, 7 *Sert-Flp^+^;Tph2^fl/fl^*; three-way repeated measures (RM) ANOVA or mixed-effects analysis followed by Tukey multiple comparisons test]. **f**, Session-by-session duration of time spent sniffing/grooming pups in the mPOA-injected virgins (top) and BNST-injected mothers (bottom). **g**, Schematic showing experimental design for the conditioned place preference test. **h**, Activation of 5HT_mPOA_ in mice with normal *Tph2* led to a significant increase in preference for the chamber that received blue light stimulation (*n* = 4 *WT*, 9 *Sert-Flp^+^;Tph2^+/+^*, 5 *Sert-Flp^+^;Tph2^fl/fl^*; two-way RM ANOVA followed by Tukey multiple comparisons test). The preference index was calculated as (time in chamber paired with blue stim – time in chamber paired with orange stim) / (total time in either chamber). **i**, Activation of 5HT_BNST_ has no significant effect in conditioned place preference (*n* = 5 *WT*, 8 *Sert-Flp^+^;Tph2^+/+^*, 7 *Sert-Flp^+^;Tph2^fl/fl^*; two-way RM ANOVA followed by Tukey multiple comparisons test). See Source Data for detailed statistics.

**Extended Data Fig. 6.**
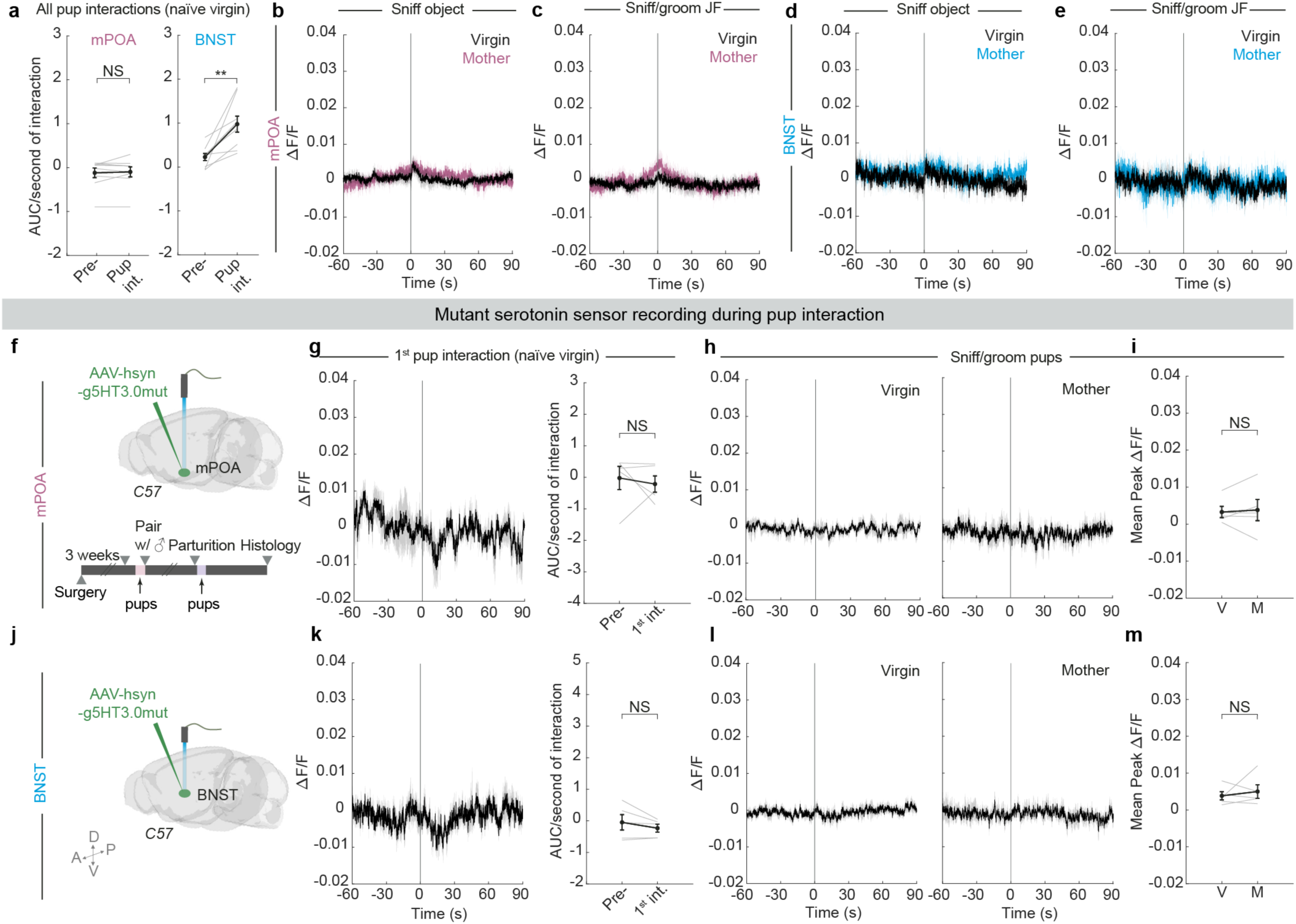
Additional characterization and controls for serotonin sensor recordings. **a,** Quantifications of area under curve (AUC) normalized to interaction duration before and during all pup interactions, including the first and subsequent interactions ranging from 9–20 trials total, in naïve virgins recorded in mPOA (left, *n* = 9) and BNST (right, *n* = 9). **b**, **c**, Peri-event time histograms (PETHs) of Δ*F*/*F* of g5-HT3.0 signal in mPOA aligned to the onset of females sniffing object (**b**) and sniffing/grooming juvenile female intruder (**c**). Each panel shows both group g5-HT3.0 signal in naïve virgins (black trace) and the same females as mothers (pink trace) (*n* = 9 virgins and mothers except in **c**, *n* = 5 for mother trace; the other 4 mothers attacked the juvenile female and were excluded as the behavior was not comparable to their non-aggressive interactions with juvenile female intruder as virgins). **d**, **e**, Same as **b**, **c** except for g5-HT3.0 imaging in BNST (*n* = 6 virgins and mothers except in **e**, *n* = 3 for mothers that had non-aggressive interactions with the juvenile female). **f**, Schematic showing the experimental setup and timeline for recording mutant serotonin sensor (g5-HT3.0mut) in mPOA. **g**, Left, group g5-HT3.0mut signal in mPOA during the first pup interaction in naïve virgins. Right, quantifications of area under curve (AUC) normalized to interaction duration before and after the first pup interaction in naïve virgins. (*n* = 5 females; Wilcoxon matched-pairs signed rank test) **h**, PETHs of Δ*F*/*F* of g5-HT3.0mut signal in mPOA aligned to the onset of females sniffing/grooming pups as virgins (left) or mothers (right). **i,** Within-animal comparisons of mean peak Δ*F*/*F* of g5HT3.0mut signal in mPOA between virgins (V) and the same animals as mothers (M) during pup interaction (*n* = 5 females; paired *t-*test). **j–m,** Same as **f–i** except for g5-HT3.0mut imaging in BNST (*n* = 5). See Source Data for detailed statistics.

**Extended Data Fig. 7.**
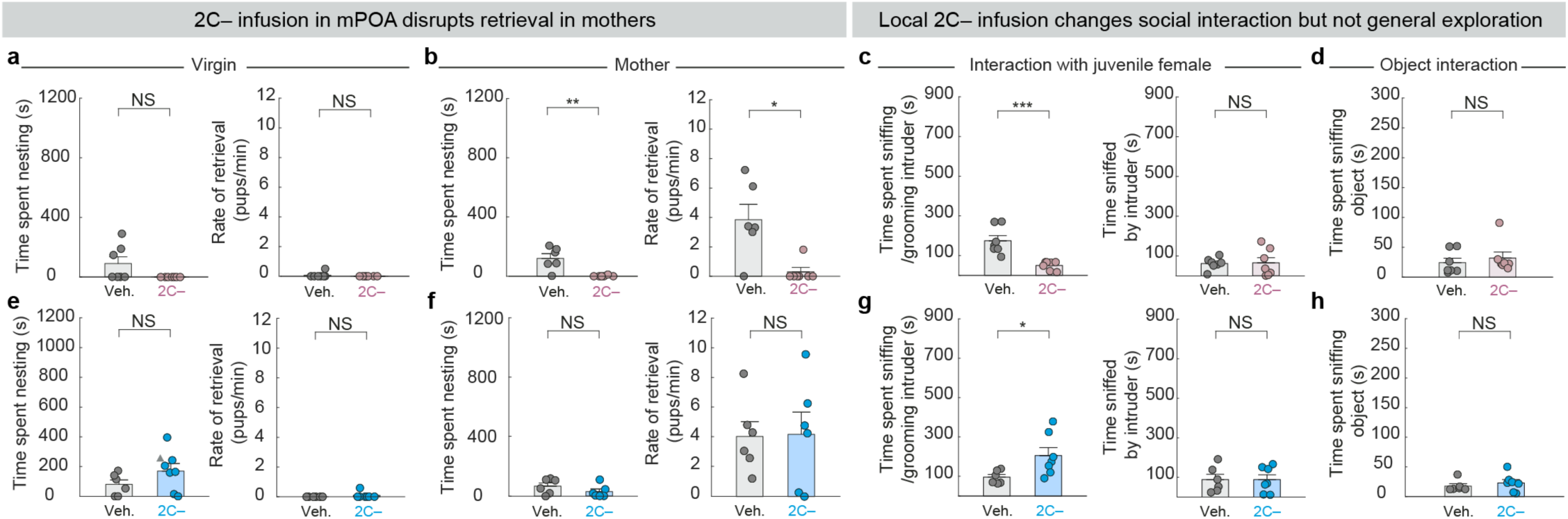
Additional behavioral characterization of local 2C– infusion. **a**, Time spent nesting around pups (left) and the rate of retrieval (right) did not differ significantly between virgin females that received 2C– in mPOA and controls (*n* = 7, 7). **b**, Females with 2C– infused in mPOA spent significantly less time nesting around pups (left) and displayed significantly lower rate of retrieval in comparison to control mice as mothers (right) (*n* = 6, 7 for mothers). **c**, Females (virgins) with 2C– infused in mPOA spent significantly less time sniffing/grooming juvenile female intruder than control mice (left). The time sniffed by the intruder does not differ between the two groups (right) (*n* = 6, 7, one control female died during labor). **d**, Object interaction does not differ significantly between virgin females that received 2C– in mPOA and controls. **e–h**, Same analyses as **a–d**, but for BNST instead of mPOA. 2C– infusion did not differ from controls except that it significantly increased time spent sniffing/grooming juvenile female intruder (**g**) (*n* = 6, 7 for virgins and *n* = 6, 6 for mothers). See Source Data for detailed statistics.

**Extended Data Fig. 8.**
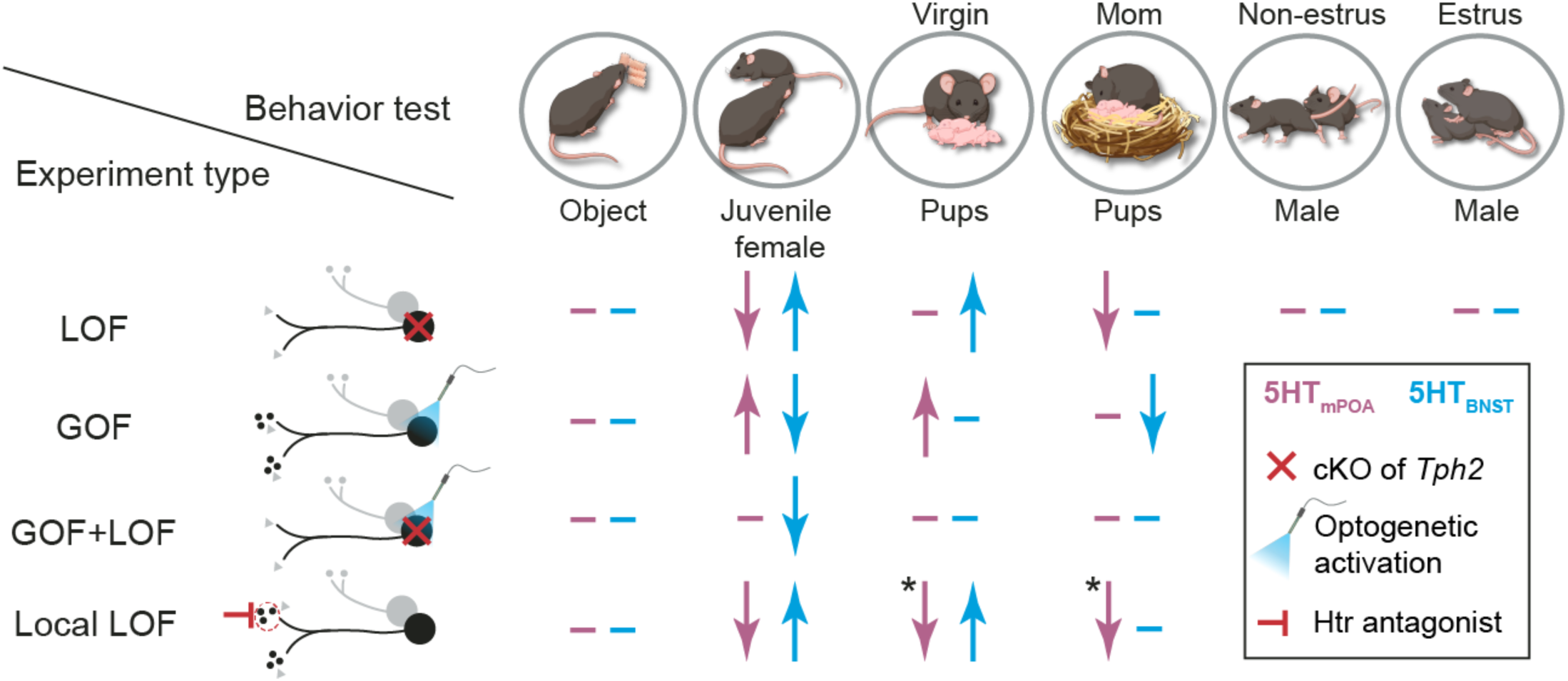
Summary of behavioral data for different loss- and gain-of-function manipulations of 5HT_mPOA_ and 5HT_BNST_. **Left,** experiment schematic. Black circles, targeted projection-specific serotonin neurons; gray circles, serotonin neurons that do not project to the target region of interest; smaller gray or black dots represent serotonin; gray triangles, other neurotransmitters or neuropeptides. **Main table,** summary of results. Pink, manipulations targeting 5HT_mPOA_; blue, manipulations targeting 5HT_BNST_; upward arrows, significantly more interaction than respective controls; downward arrows, significantly less interaction; en dash, no significant difference from controls. cKO, conditional knockout; Htr, serotonin receptor. In summary, most experiments show consistent results between the two loss-of-function (LOF) experiments—*Tph2* cKO and Htr2C– infusion—and opposite effects for between loss- and gain-of-function (GOF) manipulations. We note the following two exceptions: (I) 2C– infusion in mPOA but not 5HT_mPOA_-cKO led to lower amount of pup contact time in virgins, and (II) 2C– infusion in mPOA but not 5HT_mPOA_-cKO inhibited pup retrieval in mothers (each marked with an asterisk). These discrepancies may be due to (1) functional heterogeneity across axon collaterals in 5HT_mPOA_ and/or among serotonin receptors in mPOA; (2) the difference between long-term, gradual disruption of serotonin synthesis versus acute reduction of serotonin signaling; and/or (3) local infusion of 2C– in mPOA having a more complete coverage in reducing 5-HT activity in mPOA than retrograde Cre injection used for 5HT_mPOA_-cKO. One or a combination of these possibilities—reduced effect of potential functional heterogeneity (e.g., perturbing another target of 5HT_mPOA_ may have an opposite effect as perturbing mPOA), the lack of compensation, and higher efficiency— may have led 2C– infusion in mPOA to cause a more dramatic phenotype than 5HT_mPOA_-cKO in both virgins and mothers.

## Methods

### Animals

All experimental procedures were approved by the Stanford University Administrative Panel on Laboratory Animal Care and the Administrative Panel of Biosafety in accordance with National Institutes of Health (NIH) guidelines. All mice were housed on a 12-hour light (9 pm–9 am)/dark cycle (9 am–9 pm), with ad libitum access to food and water. *Sert-Cre* (MMRRC, Stock #017260-UCD), *Ai148* (JAX Strain#: 030328), *TRAP2* (JAX Strain#: 030323), *Sert-Flp* (JAX Strain#: 034050), *Ai14* (JAX Strain#: 7914), *Tph2^flox/flox^* (obtained from Qi Wu^41^), or wild-type C57BL6/J female mice (JAX Strain #: 000664) were used where indicated. All mice were in C57/BL6 background. Naïve virgin female mice (aged between 7–12 weeks) that had never been exposed to pups since weaning were used at the beginning of all experiments and were group-housed until surgery. After surgery, all females were single-housed until paired with a male, which was removed the day after parturition to avoid handling-induced stress near labor. All experiments were done during the dark phase.

### Virus

*AAV_retro_-hsyn-Cre-P2A-tdT*, *AAV_retro_-Ef1a-FLEx_FRT_-Cre* (denoted as *AAV_retro_-Ef1a-fDIO-Cre* on Addgene), and *AAV8-Ef1a-Con/Fon-mCherry* were purchased from Addgene. *AAV8-Ef1a-FLEx_loxP_-SOUL-P2A-tdT-WPRE- BGHpA* was custom packaged and produced by UNC NeuroTools. *AAV9-hsyn-g5-HT3.0* and *AAV9-hsyn-g5- HT3.0mut* was purchased from BrainVTA. *AAV8_retro_-hsyn-FLEx_loxP_-HA* and *AAV8_retro_-hsyn-FLEx_loxP_-mScarlet* were gifts from the lab of Xiaoke Chen. All virus titers were > 10^12^ genomic copies/mL.

### Stereotaxic surgeries

Mice were anesthetized with 1–3% isoflurane in oxygen (1 L/min) and placed into a stereotaxic apparatus (Kopf Instruments). The following coordinates (in mm) were used for virus injection: mPOA: +0.01 AP, ±0.50 ML, – 5.20 DV; BNST: –0.22 AP, ±0.85 ML, –3.99 DV; DR: –4.3 AP, 1.10 ML, –2.85 DV, with 20° ML angle; MR: – 4.20 AP, 1.15 ML, –4.50 DV, with 14° ML angle (AP is relative to bregma as defined by the Franklin and Paxinos mouse brain atlas^54^). Virus was delivered to the site of injection via a glass capillary at a rate of 100 nL/min. After surgery, mice recovered on a heating pad until ambulatory.

To perform Ca^2+^ imaging on serotonin neurons in freely moving female mice, a zirconia fiber optic cannula (Doric Lenses, 0.66 NA, 400 μm, borosilicate, flat tip) was implanted over either the DR (–4.3 AP, 1.10 ML, 20° ML angle) or MR (–5.22 AP, 0.00 ML, 14° AP angle), and was secured to the skull using dental cement (Parkell, C&B metabond, S371, 396, 398). Implanted mice were single-housed until imaging at least two weeks later.

To map the axon projections of serotonin neurons during pup interaction or home cage activity, *AAV8-Ef1a- Con/Fon-mCherry* was injected into both DR and MR [three injections at each site (200 nL, 200 nL, 100 nL; 0.05 mm apart along the DV injection trajectory for a total of 500 nL)] of *TRAP2;Sert-Flp* naïve virgin females. Two weeks later, the injected mice were randomly assigned to be TRAPed during either pup interaction or home cage activity.

To perform retrograde labeling of mPOA- and BNST-projecting serotonin neurons, 250 nL of *AAV8_retro_-hsyn- FLEx_loxP_-HA* was injected bilaterally into either mPOA or BNST of *Sert-Cre* females. To examine the overlap between mPOA- and BNST-projecting serotonin neurons, and the overlap targeting same population between two separate injections, *AAV8_retro_-hsyn-FLEx_loxP_-HA* and *AAV8_retro_-hsyn-FLEx_loxP_-mScarlet* (250 nL) were injected bilaterally into either different sites (mPOA, BNST) or the same sites sequentially (mPOA, mPOA). The histology was examined three weeks after the injection.

To delete *Tph2* from projection-defined serotonin neurons, *AAV_retro_-hsyn-Cre-P2A-tdT* was injected bilaterally into mPOA (300 nL per hemisphere) or BNST (300 nL per hemisphere) of *Tph2^flox/flox^*females. For control, the same virus was injected into the same regions of *WT* mice. Naïve virgin female behavioral experiments were performed 17 days post *AAV_retro_-hsyn-Cre-P2A-tdT* injection for *Tph2* conditional knockout experiments.

To optogenetically activate mPOA- or BNST-projecting serotonin neurons, 300 nL *AAV_retro_-Ef1a-FLEx_FRT_- Cre* was injected bilaterally into mPOA or BNST, and 400 nL of *AAV8-Ef1a-FLEx_loxP_-SOUL-P2A-tdT-WPRE- BGHpA* was injected respectively into DR and MR. The cannula (Thorlabs, CFML12L02, 1.25 mm SS Ferrule, 200 µm Core, 0.39 NA) was inserted at –6.50 AP, 0.00 ML into the brain at 30° AP angle such that the tip of the cannula sat just under the *brain* surface to allow the cone of light to reach both DR and MR, taking advantage of the ultra-sensitivity of SOUL. The cannula was secured to the skull using dental cement (Parkell, C&B Metabond). Naïve virgin female behavioral experiments were performed three weeks after the injection.

To record serotonin activity in mPOA or BNST, 300 nL *AAV9-hsyn-g5-HT3.0* was injected unilaterally into mPOA or BNST. A zirconia fiber optic cannula (Doric Lenses, 0.66 NA, 400 μm, borosilicate, flat tip) was then implanted over mPOA or BNST and secured to the skull using dental cement (Parkell, C&B Metabond). Mice were imaged using the fiber photometry system three weeks after virus injection.

For drug infusion, a double-lumen, 26-gauge threaded guide cannula was implanted such that the end of the infusion cannula sits at the top of the targeted infusion site. For mPOA, the guide cannula dimensions were with 1.0-mm separation and cut 5 mm below the pedestal. For BNST, the guide cannula dimensions were with 1.5-mm separation and cut 5 mm below the pedestal. The guide cannula was secured to the skull using dental cement (Parkell, C&B Metabond). A bilateral dummy cannula was inserted into the cannula guide to prevent blockages before drug infusion and seal the cannula after infusion. The top of the guide cannula was covered with a plastic screw cap to keep the cannula clean and protected from animal activities such as allogrooming and chewing. Mice were used for infusion experiments at least 10 days after the implant.

### Fiber photometry

Each mouse was connected by the implanted cannula via a ceramic split mating sleeve to a 0.57-NA, 400-μm low autofluorescence optical fiber patch cord, which then connects to a custom photometry acquisition system through a pigtailed fiber optic rotary joint (0.57-NA, 400-μm). All animals were habituated to the fiber connection and were imaged using modulated 405 nm and 490 nm LEDs (Thorlabs, M405F1 and M490F3). The emitted fluorescence signal passed back via the same fiber optics through the GFP emission filter (Thorlabs, MF525-39), and was detected by a Newport 2151 photoreceiver. Fluorescence and behavioral data were synchronized and collected through an RZ5 real-time processor (Tucker-Davis Technologies, TDT) at a sampling frequency of 1.017e03 Hz for fiber photometry data and 20 frames per second (fps) for behavioral data using the Synapse software (TDT). Fiber photometry data were collected during freely moving interactions respectively with a juvenile female intruder, pups, dummy pups (made of pink PIG® cloth), and a sexually experienced male. Due to the variability in natural behaviors, we prioritized collecting between 9–20 trials from every animal for most behaviors, rather than restricting our recording sessions to the same duration for all animals. We note that similar to our observation in the pup-TRAP experiment, some virgin females retrieved or re-nested around pups after approximately an hour of pup exposure. The recording used for naïve virgin pup interaction analysis was restricted to the period before any sign of pup sensitization (e.g., retrieval or pup-oriented nesting behavior).

Behavioral data were annotated by experimenters blind to neural responses prior to the calculation of ΔF/F using the same definitions delineated in the ‘**Behavioral assays and analyses**’ section below. Fiber photometry data were analyzed in MATLAB. The first 5 to 10 seconds were removed from all demodulated streams to exclude the artifact associated with the onset of LEDs turning on. Each channel was then downsampled to 101 Hz. The 405-nm stream was fitted to the 490-nm stream using a least-squares fit, with ΔF/F computed as (F490 – fittedF405)/fittedF405. The ΔF/F was then aligned to the onset of each trial of a particular behavior and averaged across all trials for each animal. The PETHs were then made after averaging across recorded animals. Fiber optic placement was verified for all mice by histology.

### TRAP

4-hydroxytamoxifen (4OHT; Sigma, Cat# H6278) was dissolved in ethanol (20 mg/mL) by shaking at 37°C for 40 min, aliquoted and then stored at –20°C. On the day of TRAP experiments, a 1:4 mixture of castor oil:sunflower seed oil (Sigma, Cat # 259853 and S5007) was added to 4-OHT aliquot that had been first shaken at 37°C for 30 min. The 4-OHT mix was then vortexed and then centrifuged under vacuum until all ethanol evaporated, leaving the final concentration at 10 mg/mL. All injections were delivered intraperitoneally (i.p.) at 50 mg/kg, an hour after pup interactions or home cage activity. The animals were placed back to their home cage after the injection and allowed to continue pup interaction or home cage activity for another three hours. Note that pup-TRAP and HC-TRAP were always performed together on the same day at the same time, in separate sound-proof chambers in the same behavior room, and injected with the same 4-OHT aliquot prepared fresh on the same morning.

### Whole brain clearing

At least five weeks after TRAP, mice were perfused transcardially with a phosphate-buffered saline (PBS) and ice-cold 4% paraformaldehyde (PFA). The dissected brains were post-fixed in 4% PFA on a shaker at 4℃ overnight. On the next day, samples were washed three times with 1×PBS containing 0.02% NaN₃ at room temperature, with gentle shaking for one hour between each wash. All samples were then processed in 5 mL volumes based on the previously described modified AdipoClear immunolabeling protocol^32^. Briefly, the samples were dehydrated with MeOH gradient in B1n (a mixture of distilled H_2_O, 10% Triton-X, Glycine, 10N NaOH and 5% NaN_3_) (20%, 40%, 60%, and 80%) before two washes in 100% MeOH. The samples were then incubated overnight in 2:1 dichloromethane (DCM):MeOH mixture overnight. On the following day, the samples were washed in 100% DCM and two times in 100% MeOH before bleaching in 5:1 MeOH:30% H_2_O_2_ mixture. Following a rehydration step with reversed MeOH/B1n gradient, the samples were washed in B1n, two times in 5% DMSO/0.3M Glycine/PTxwH (a mixture of distilled H_2_O, 10× PBS, 10% Triton-X, Tween 20, Heparin and 5% NaN_3_) and the three times in PTxwH. For immunostaining, whole brains were incubated with primary antibody (rabbit anti-RFP, Rockland, 600-401-28379, 1:500) for 11 days and later secondary antibody (donkey anti-rabbit AlexaFluor 647, ThermoFisher Scientific, 1:500) for 7 days, both steps on the shaker at 37℃. At the end of each antibody step, the samples were washed five times in PTxwH and then one wash per day for two additional days. For tissue clearing, the samples were dehydrated with MeOH gradient in water, washed three times in 100% MeOH, incubated in 2:1 DCM:MeOH mixture overnight at room temperature, washed in 100% DCM twice the following day and transferred to be cleared in dibenzyl ether. The cleared samples were then transferred to ethyl cinnamate, washed at least once per day for three additional days before imaging.

### Light-sheet imaging and analysis

Cleared brains were imaged using a light-sheet microscope (Blaze, Miltenyi Biotec) in ethyl cinnamate (Millipore Sigma, 112372) at 1× magnification using 647-nm imaging laser with dynamic focusing at a z-step size of 3 μm to capture stained axons at a final voxel size was 5.9 × 5.9 × 3 μm. Each brain was also imaged using the 488-nm laser but without dynamic focusing to obtain the sample autofluorescence. Axons were then segmented from the raw 647-nm image stack using TrailMap^32^. The probabilistic volume of segmented axons was then binarized at eight thresholds from 0.2 to 0.9, skeletonized, and then summed together with each weighted at the initial probability threshold. The 488-nm autofluorescence channel was used for registration, where the image stack was aligned to the Allen Institute’s 25-μm reference brain generated by serial two-photon tomography using linear and non-linear transformations from the Elastix toolbox^55,56^. The same parameters were then applied to warp the axon volume into the same Allen Institute’s Common Coordinate Framework (CCF)^57^ space. In consistency with a previous study^18^, regions defined exclusively by layers were collapsed into the parent region such that layers 1– 6 of prelimbic area, for instance, are jointly labeled as “prelimbic area”. The Allen brain atlas mask was then used to quantify axon on a region-by-region basis. Axon labeling density was computed as the total number of voxels classified as axon-positive divided by the region volume. The normalized labeling density was computed by range normalizing the axon labeling density such that it ranged from 0–1 for every brain across the two TRAP conditions. Images were generated using Fiji and Imaris software.

### Behavioral assays and analyses

#### Interactions with juvenile female

A novel juvenile female (JF) (21–25 days old) was introduced to the home cage of the test mouse for 15 min of free interactions. Following this, the JF was removed from the test mouse’s home cage. ‘Sniff/groom juvenile female intruder’ was defined as when the test mouse’s snout made contact with the juvenile female body and included both anogenital sniffing and facial grooming. ‘Sniffed/groomed by juvenile female intruder’ was defined as when the juvenile female initiated snout contact with the test mouse’s body. ‘Aggression’ was defined as when physical attacks including biting, episodes of wrestling, and chasing occurred.

#### Object interactions

A novel plastic object was placed on the opposite side of the nest inside the test mouse’s home cage. Free interactions were recorded for a total of 5 min. Novel object interactions were defined as the amount of time the test mouse extended her body to sniff the object or when her snout made direct contact with the novel object. Locomotion was measured during the same object interaction session by tracking key body parts of the test animal using methods described in the ‘**Animal pose estimation**’ section below.

#### Pup interactions

To assess pup interaction in naïve virgin females, four pups (at age of P1–P4) were placed on the opposite side of the experimental mouse’s nest in the home cage, and their free interactions were recorded for 20 minutes. To induce pregnancy, each virgin female was paired with a male. Between one to two days after parturition, the male was removed from the cage. Four pups were removed from the nest, kept separate from the mother briefly, and then placed back to the cage on the opposite side of the nest. Free interactions were recorded for 20 minutes. ‘Pup sniffing/grooming’ was defined as when the test mouse’s snout was in direct contact and engaging in close interactions with the pups. ‘Huddling’ was defined when the test mouse made contact with the pup with its body centroid above the pups without engaging the snout. ‘Pup contact’ was the sum of both pup sniffing/grooming and huddling. ‘Nesting’ was defined as when the test mouse picked up nesting materials with her mouth and deposited them around or on the pups. ‘Retrieval’ was defined as when the test mouse picked up the pups with her mouth and carried the pups back to the nest. The rate of retrieval was calculated as the number of pups retrieved per minute.

#### Sexual interactions

A sexually experienced adult male was placed into the home cage of the test female. The female’s reaction was examined when the male mounted, which was defined as when the male moved on top of the female, clasped his forepaws around the female and began pelvic thrust. ‘Reject mating’ was aligned to when the female escaped from the mounting male. ‘Mating’ was defined as when the female remained beneath the male during intromits and was aligned to the first intromission. Receptivity index was calculated as the ratio of male mounts that resulted in mating. The estrus cycle of the female was determined by vaginal cytology^58^. Females were considered to be in estrus when the main cell type was cornified epithelial cells, and non-estrus if primarily leukocytes.

All behaviors were recorded in the home cage of the test mouse, which was placed inside a dark behavior chamber with only infrared illuminators. The behavior chamber was housed inside a room illuminated by only red light. All behaviors were recorded by a camera located above the home cage, controlled by Synapse software (TDT) at 20 fps. The infrared filter was removed from the Logitech camera to capture videos under only infrared illumination. Behavioral videos were manually annotated by experimenters blind to the test condition using Boris software^27^.

### Animal pose estimation

Animal pose estimation was performed using Lightning Pose (version 1.6.1)^59^ to generate a single animal 7 body part model (body parts: nose, left ear, right ear, centroid, left hip, right hip, and tail base). To create the training, testing and validation sets, 870 video frames taken from 282 videos and 92 animals were manually labelled, with 80% used for training, 10% for testing, and 10% for validation. Training and testing were conducted using a ResNet101-based neural network pretrained on ImageNet data with a DeepLabCut image augmentation pipeline (DLC imgaug), an Adam optimizer, and an initial learning rate of .001, and loss calculated as mean squared error. The model was trained for up to 750 epochs or until plateau^60^, and was validated with 5 shuffles and temporal and single view principal component-based (pca_singleview) losses. For an image of size 256 by 256, the average pixel error was 29.92 pixels for test, 29.87 pixels for train, and 29.92 pixels for validation. Average pca singleview reprojection error was: 13.06 pixels for test, 6.74 pixels for train, and 12.04 pixels for validation. A probability cutoff of .9 was used to condition future predicted coordinates. This network was then used to analyze videos from similar experimental settings. Following predictions on all frames from all videos, custom code was used to convert tracking data to a format compatible with the Python package Ensemble Kalman Smoothing (EKS), which was then used to ensemble and smooth pose estimates to improve tracking^59^. The frame-by-frame nose position obtained from the automatic pose tracking was then used to calculate total distance travelled and construct heat maps showing animal localization throughout the behavioral session.

### Optogenetic activation

Blue (473 nm, SLOC Lasers) and orange (589 nm, OEM Laser Systems) lasers were coupled to the RZ5 processor and controlled by Synapse software (TDT). Each stimulation was programmed to be 5 Hz for a total duration of 30 s. The laser power at the end of the cannula tip was calibrated to be 14–15 mW. At the beginning of the experiment, mice were connected to the lasers through a ferrule patch cable (Thorlabs, M83L01, 200 µm core, 0.39 NA, FC/PC to Ø1.25 mm) via a ceramic split mating sleeve. After 5 minutes of habituation, one of the three types of stimuli tested (four pups aged between P1–P4, a juvenile female or an object) was placed on the opposite side of the nest in the home cage. The test mouse was first allowed to interact with the stimulus for 5 minutes to attenuate novelty-driven exploration. The stimulation then alternated between blue and orange laser every 5 minutes for a total of 20 minutes. The mice were randomly assigned to receive blue or orange stimulation first. All interactions were recorded in the home cage.

### Conditioned place preference test

The conditioned place preference (CPP) apparatus consisted of a custom-built apparatus with three separate chambers: two conditioning chambers with distinct visual cues (striped or dotted floor) and a neutral middle chamber lacking distinguishing features. All experiments were conducted with LED illumination at the top of the chamber to facilitate feature detection. During the pretest, mice were placed in the neutral middle chamber with no restraints and allowed to explore the entire apparatus freely for 10 minutes. Time spent in each chamber was recorded to assess baseline preference. After the pretest, mice received a 30-second session of optogenetic stimulation (473-nm blue or 590-nm orange laser, randomized across animals) in the home cage, and were then placed and confined in chamber 1 with striped flooring for 15 minutes. After this first conditioning session, mice were returned to their home cage and stimulated with the alternate wavelength (orange or blue) for 30 seconds. They were then placed and confined in chamber 2 with dotted flooring for an additional 15 minutes of conditioning. The order of light pairing (laser wavelength to chamber) was counterbalanced across animals. Mice were returned to their home cage at the end of conditioning. Finally, during post-test, mice were again placed in the neutral middle chamber with free access to both chambers for 10 minutes. Time spent in each chamber was recorded to assess preference in comparison to the pretest. The preference index was calculated as (time in chamber paired with blue stim – time in chamber paired with orange stim) / (total time in either chamber).

### Immunohistochemistry and imaging

Mice were anesthetized with 2.5% avertin and perfused using phosphate buffered saline (PBS) followed by 4% ice-cold paraformaldehyde (PFA). The dissected brains were post-fixed in 4% PFA overnight at 4℃ and then switched to 30% sucrose for 24–48 hours at 4℃. Brains were flash frozen in Optimum Cutting Temperature (OCT), sectioned at 60 µm using a cryostat (Leica CM1850), immunolabeled and stained with DAPI (1:1000) using previously published procedures^18,48^. Free-floating sections were incubated with primary antibodies diluted in 0.1% PBST overnight at 4℃ at the following concentrations: Goat anti-Tph2 (abcam, ab121013, 1:1000), Rabbit anti-RFP (Rockland, 600-401-379, 1:1000), Chicken anti-GFP (abcam, ab13970, 1:1000), Guinea pig anti-Cre-recombinase (Synaptic Systems, 257005, 1:1000), Guinea pig anti-Fos (Synaptic Systems, 226308, 1:1000; used when another rabbit primary antibody was used simultaneously), Rabbit anti-c-fos (Synaptic Systems, 226008), Rabbit anti-DOPA Decarboxylase (also known as aromatic L-amino acid decarboxylase or AADC; Thermo Fisher, PA1-4651, 1:250), Rabbit anti-HA-Tag (Cell Signaling Technology, 3274, 1:500), Rat anti-mCherry (Thermo Fisher, M11217, 1:500). Following three 1×PBS washes the next day, the sections were incubated in the corresponding secondary antibodies (1:1000 for all) including: Donkey anti-goat, Alexa 647 conjugate (Jackson ImmunoResearch, 705-605-147), Donkey anti-rabbit, Cy3 conjugate (Jackson ImmunoResearch, 711-165-152), Donkey anti-rabbit, Alexa 488 conjugate (Jackson ImmunoResearch, 711-545- 152), Donkey anti-chicken, Alexa 488 conjugate (Jackson ImmunoResearch, 703-545-155), Donkey anti-guinea pig, Cy3 conjugate (Jackson ImmunoResearch, 706-165-148), Donkey anti-guinea pig, Alexa 488 conjugate (Jackson ImmunoResearch, 706-545-148), Donkey anti-rat, Cy3 conjugate (Jackson ImmunoResearch, 712-165- 150).

For serotonin staining, mice were perfused as described above. The dissected brains were post-fixed in 4% PFA overnight at 4℃, and then sectioned at 60 µm with a vibratome (Leica VT1000S). The sections were first processed with an antigen retrieval step, where sections were heated in 1× citrate buffer (pH6.0; diluted from 10× citrate buffer made with 82 mL 1M sodium citrate and 18 mL 1M citric acid in water for a total volume of 1 L) until boiling. The sections were then incubated with Goat anti-5HT polyclonal antibody (abcam, ab66047, 1:1000) on a rocker at room temperature overnight. Following three 1×PBS (5 min) washes the next day, the sections were incubated with Donkey anti-goat, Cy3 conjugate (Jackson ImmunoResearch, 705-165-147, 1:1000) and DAPI (1:1000) at room temperature overnight.

Processed sections were mounted, coverslipped with Fluoromount-G (SouthernBiotech, 0100-01) and then imaged with Zeiss LSM 900 confocal microscope.

### Drug microinfusion

Serotonin receptor antagonist (SB242084) (Tocris Biosciences, 2901) was dissolved in DMSO to make a stock solution at the concentration of 50 µg/µL. On the day of behavioral experiments, the stock solution was further diluted in 0.9% saline to make a working solution at 2 µg/µL. To reduce handling stress during the infusion process, mice were anesthetized with 1–2% isoflurane in oxygen (1 L/min) and placed into a stereotaxic apparatus. For mPOA infusion, an internal cannula with a 0.5-mm protrusion beyond the guide cannula was used such that the infusion tip reached the top quarter of mPOA. For BNST, an internal cannula with no protrusion (0.0 mm) beyond the guide cannula was used. Drugs were then manually infused through the internal cannula using a 10- µL syringe (Hamilton, 80383) at a rate of ∼350 nL/min to deliver 1.5 µg of drug. Infusions were performed bilaterally. After infusion at each site, the internal cannula was held in place for 2 minutes to reduce efflux. Following the removal of the internal cannula, the dummy cannula was reinserted into the cannula guide and the implant was resealed with a plastic cap. Mice were allowed to recover in the home cage for 17 min before behavioral tests.

### Statistics

All statistical analyses were performed using MATLAB and Prism software. Data were shown as means ± SEMs. For data that passed Shapiro–Wilk test for normality, parametric tests were used. Differences between two samples were analyzed using two-tailed unpaired *t*-test or paired *t*-test. Group differences were analyzed using one-way, two-way, or three-way analysis of variance (ANOVA) depending on the experimental design or mixed-effects analysis, followed by Tukey or Šídák’s multiple comparisons test. For paired virgin-mother data sets that contain missing values due to random death during labor, mixed-effects analyses were used. For data that did not pass normality tests, nonparametric tests including Mann-Whitney U-test were used. Sample sizes were selected based on previous papers. Unless specified otherwise, all statistical analyses were two-tailed, and significance was defined as *p* < 0.05. All experiments consisted of 2–4 cohorts that were analyzed together at the end. Full details of statistical test, sample sizes and statistical results are reported for each panel of each figure in the Source Data.

## Acknowledgements

We thank Yuan Yuan and Sijia Fan from Xiaoke Chen Lab for viruses and advice on drug infusion; Daniel Pederick and Chung-ha Oh Davis for their advice on behavioral experiments; Jing Ren for advice on fiber photometry; Nirao Shah, Jing Ren, and all members of the Luo Lab, especially Tom Hindmarsh Sten, Jalal Baruni, Colleen McLaughlin, Chloe Bair Marshall, Airi Yoshimoto, Lijun Qi, Jun Song, and Ying Hu for advice and feedback on the manuscript. We thank Jing Ren, Jun Song, Yunming Wu, and Jalal Baruni for sharing mice and reagents; Yunming Wu for the antigen retrieval protocol; Kenneth Magpusao and Carlota Manalac for assistance caring for the mouse colony; Mary Molacavage and David Luginbuhl for help with project logistics. This work was supported by National Institutes of Health grant (1R01NS131987-01 to L.L. and S.W.L.). L.L. is an investigator of Howard Hughes Medical Institute.

## Author Contributions

L.L. and S.A.X. conceived the project and designed the experiments. L.L. supervised the project. S.A.X. performed most of the experiments and data analyses, and prepared figures. C.C.C. performed the conditioned place preference experiment, and helped perform other optogenetics-related behavioral and histological experiments. C.C.C. and V.M.C. helped run drug infusion experiments and collect histology. P.H. and C.D. contributed to experiments examining the local action of serotonin on mPOA neurons and provided critical feedback on the experiments. V.T., D.B., and S.W.L. performed animal pose estimation. F.D. and Y.L. shared the latest serotonin sensor, g5-HT3.0. S.A.X. and L.L. wrote the manuscript with feedback from all coauthors.

## Competing interests

The authors declare that they have no competing interests.

## Data availability

Source data are provided in the manuscript. All raw fiber photometry and behavioral recordings are deposited in the nwb format (Neurodata Without Borders) on DANDI: https://dandiarchive.org/dandiset/000892 (access will be unembargoed upon publication).

## Code availability

Code is available on github: https://github.com/vtsai881/Xiao_et_al_2025

## Supplementary Materials

**Video 1** | Coronal map showing the voxel-wise difference in average axon labeling density between the pup- TRAPed and HC-TRAPed brains (related to Fig. 2g). Darker red indicates higher mean axon density in the pup- TRAP condition.

**Video 2** | A *WT* control female interacted minimally with pups as a naïve virgin, but spent extensive time sniffing, grooming and maintaining physical contact with pups after parturition.

**Video 3** | Three example 5HT_mPOA_-cKO mothers interacted minimally with pups, despite normal pup retrieval.

**Video 4** | Three example 5HT_BNST_-cKO pup-naïve virgin females spent extensive time sniffing, grooming and maintaining physical contact with pups. Note that the first two females dragged nesting materials to re-nest around the pups, a behavior exhibited by 7 out of 9 females examined. The remaining two 5HT_BNST_-cKO females retrieved pups to their nest instead of relocating their nest completely, one of which is shown in the third example.

## Notes

### Competing Interest Statement

The authors have declared no competing interest.

